# Recurrent Integration of Human Papillomavirus Genomes at Transcriptional Regulatory Hubs

**DOI:** 10.1101/2021.08.30.457540

**Authors:** Alix Warburton, Tovah E. Markowitz, Joshua P. Katz, James M. Pipas, Alison A. McBride

## Abstract

Oncogenic human papillomavirus (HPV) genomes are often integrated into host chromosomes in HPV-associated cancers. HPV genomes are integrated either as a single copy, or as tandem repeats of viral DNA interspersed with, or without, host DNA. Integration occurs frequently in common fragile sites susceptible to tandem repeat formation, and the flanking or interspersed host DNA often contains transcriptional enhancer elements. When co-amplified with the viral genome, these enhancers can form super-enhancer-like elements that drive high viral oncogene expression. Here, we compiled highly curated datasets of HPV integration sites in cervical (CESC) and head and neck squamous cell carcinoma (HNSCC) cancers and assessed the number of breakpoints, viral transcriptional activity, and host genome copy number at each insertion site. Tumors frequently contained multiple distinct HPV integration sites, but often only one “driver” site that expressed viral RNA. Since common fragile sites and active enhancer elements are cell-type specific, we mapped these regions in cervical cell lines using FANCD2 and Brd4/H3K27ac ChIP-seq, respectively. Large enhancer clusters, or super-enhancers, were also defined using the Brd4/H3K27ac ChIP-seq dataset. HPV integration breakpoints were enriched at both FANCD2-associated fragile sites, and enhancer-rich regions, and frequently showed adjacent focal DNA amplification in CESC samples. We identified recurrent integration “hotspots” that were enriched for super-enhancers, some of which function as regulatory hubs for cell-identity genes. We propose that during persistent infection, extrachromosomal HPV minichromosomes associate with these transcriptional epicenters, and accidental integration could promote viral oncogene expression and carcinogenesis.

## INTRODUCTION

Persistent infection with high-oncogenic risk HPV types is responsible for almost all cervical and ∼70% oropharyngeal carcinomas ^1^. One factor that can contribute to oncogenic progression of HPV-positive lesions is integration of the viral genome into host chromatin. Integration is associated with increased genetic instability in high-grade cervical intraepithelial neoplasia (CIN) and cervical and oropharyngeal carcinomas ^2–9^ due to dysregulated expression of the viral oncoproteins, E6 and E7. Many studies have compared the human genomic regions associated with HPV integration sites to elucidate the mechanisms that might promote integration and carcinogenesis. Here, we have curated datasets from The Cancer Genome Atlas and other published sources to define a rigorous database of integration breakpoints and correlated these with “in-house” datasets of common fragile sites and enhancer elements defined in cervical carcinoma cells.

Integration of HPV DNA occurs in all human chromosomes; however, integration sites are often found within or in close proximity to common fragile sites ^10–13^. Common fragile sites are regions of the genome that have difficulty completing replication and, as such, are susceptible to chromosome breakage in mitosis. They are prone to replication stress that can be due to a shortage of replication origins or clashes between replication and transcriptional processes. Therefore, they vary in fragility depending on cell type and disease state ^14^. Most previous studies that documented an association of HPV integration sites with common fragile sites used the classical FRA-regions that were defined cytogenetically in lymphocytes ^9–13,15^. As such, these fragile sites are not cell-type specific and are often large, poorly defined regions that cover a large proportion of the human genome. FANCD2 is required for resolution of these genetically unstable sites and, as such, is a marker of common fragile sites ^16–18^.

HPV integration sites occur frequently in amplified regions of the host genome, and focal amplification of cellular flanking sequences at sites of viral integration are frequently observed in HPV-positive tumors ^6,7,9,15^. Co-amplification of the viral genome and flanking cellular sequences can result from unlicensed initiation of replication at the viral origin resulting in endoreduplication ^19–21^. Subsequent recombination can result in amplified tandem repeats. Genome amplification can also occur at common fragile sites by breakage-fusion-bridge cycles ^22^.

HPV integration is also enriched at transcriptionally active regions of the host genome ^15,23^. We previously identified an HPV16 integration site in the W12 20861 cervical cell line that was adjacent to a cell-type specific enhancer. Co-amplification of this regulatory element and the viral genome to approximately 25 copies resulted in the formation of a super-enhancer-like element to drive high viral oncogene expression ^24,25^. This ‘enhancer-hijacking’ is a novel mechanism by which HPV integration can promote oncogenesis; however, it is unclear how common this mechanism is in HPV-associated cancers.

The aim of this study was to examine the association among HPV integration loci, common fragile sites, and genome amplification to determine whether insertion of HPV genomes adjacent to active cellular enhancers often resulted in viral ‘enhancer-hijacking’, and whether genetic instability could result in co-amplification of viral-cellular regulatory repeats to drive oncogenic progression of HPV-associated cancers. We have extended our previously published work ^26^ to generate a common fragile site dataset in cervical carcinoma cell lines using higher resolution mapping of FANCD2 binding and have mapped cellular enhancers and super-enhancers in an HPV16-positive cell line derived from a cervical lesion^27^ using H3K27ac and Brd4 ChIP-seq (**Figure 1**). Here we compare HPV integration sites with these cervical cell-type specific enhancer and fragile site datasets.

**Figure 1.**
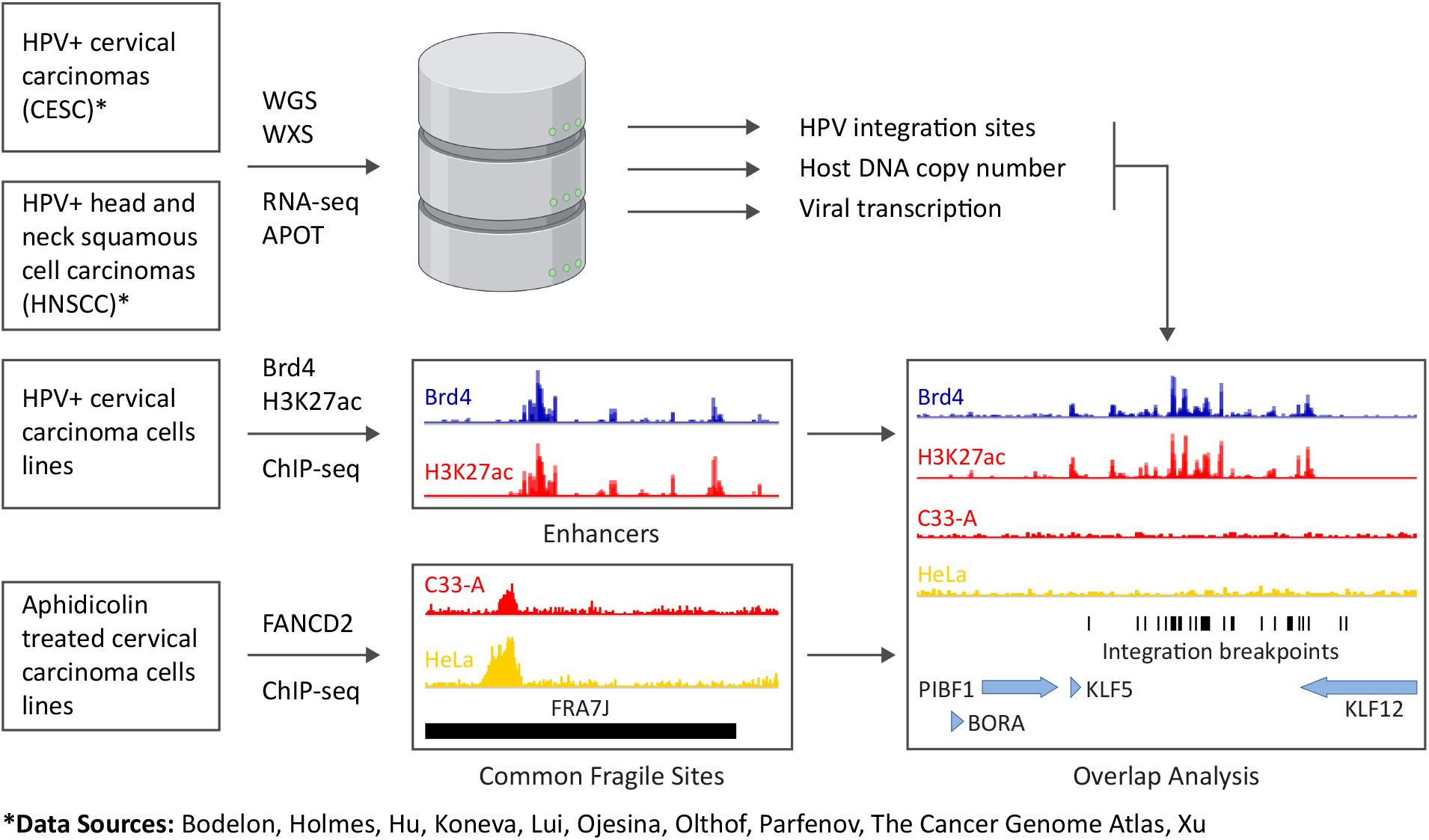
Overview of datasets. Schematic representation of datasets used for overlap analysis of CESC and HNSCC integration sites with enhancers mapped in W12 cervical keratinocytes and FANCD2-associated common fragile sites mapped in C33-A and HeLa cervical carcinoma cell lines. *APOT, Amplification of papillomavirus oncogene transcripts; WGS, Whole genome sequencing; WXS, Whole exome sequencing*.

## RESULTS

### CESC and HNSCC tumors frequently contain multiple, clustered HPV integration breakpoints

A dataset of HPV integration breakpoints was assembled from various sources ^5,6,8,9,28–35^, as outlined in **Figure 1** and **Supplementary Table S1**. Integration breakpoints were defined as the junctions between the viral and host chimeric reads within the human reference genome. A total of 1,299 integration breakpoints from 333 cervical carcinomas (CESC) and 119 integration breakpoints from 41 head and neck squamous cell carcinomas (HNSCC) were included in this study (**Supplementary Figure S1** and **Supplementary Table S2-S3)**. We found that many tumor samples contained multiple integration breakpoints that could have resulted from either independent integration events at different chromosomal loci or from amplification of a single integration site resulting in a cluster of multiple, closely spaced breakpoints. To classify this, we defined an integration locus as either a single HPV insertion breakpoint, or as multiple, closely spaced breakpoints (a cluster), **Figure 2a**. Clustered breakpoints within the same chromosome that had a maximum distance of 3 Mb between the most 5’ and 3’ breakpoints were classified as a single integration locus. Based on this classification, the total number of integration loci analyzed in our study was 584 for CESC samples and 58 for HNSCC samples. Tumors with multiple integration loci were observed in 28.8% of CESC and 22.0% of HNSCC tumors.

**Figure 2.**
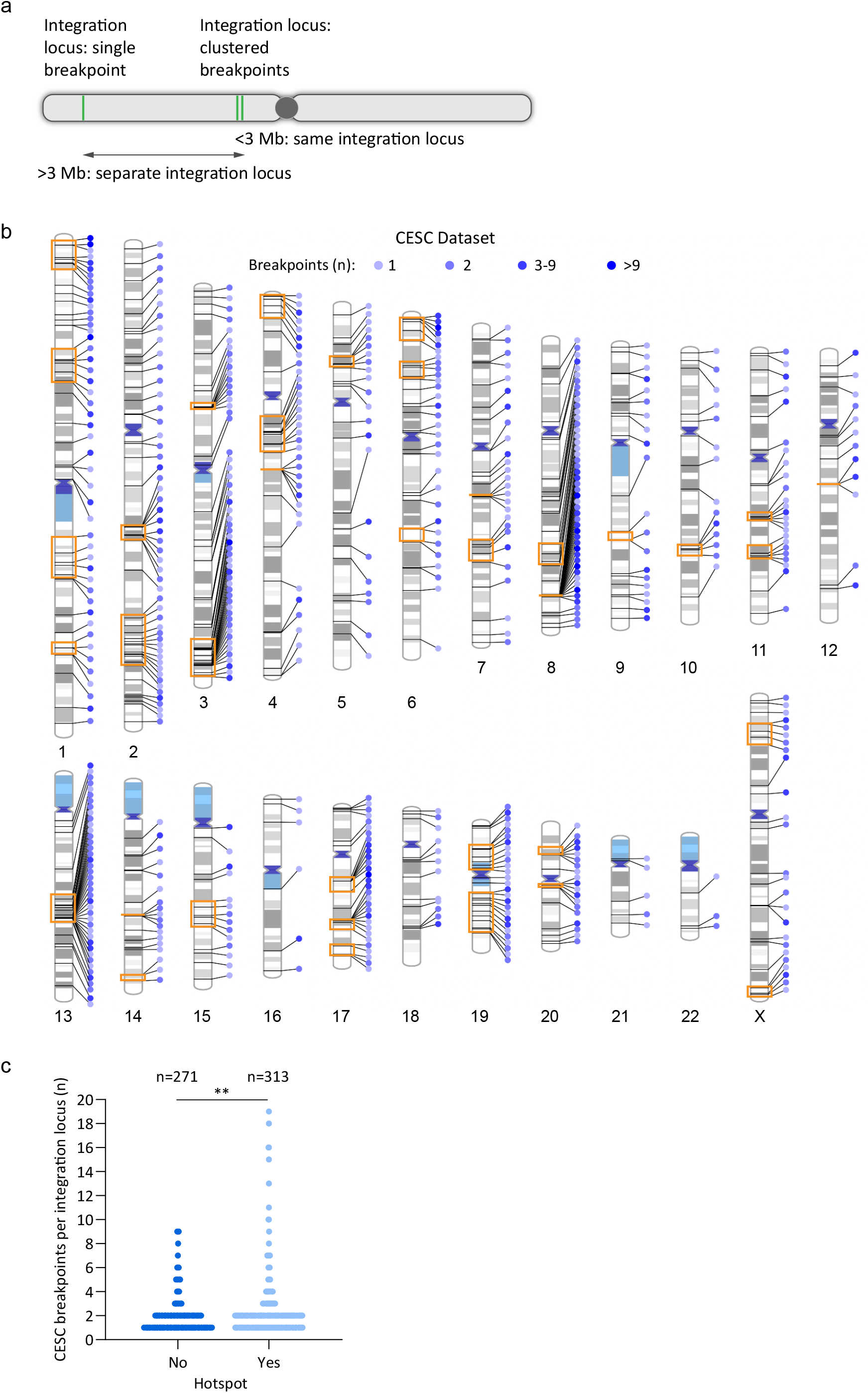
HPV integration loci frequently contain clustered insertional breakpoints. (**a**), Schematic representation of HPV integration breakpoints and loci. Green lines represent integration breakpoints. Integration loci are defined as either a single breakpoint, or multiple, closely spaced breakpoints (a cluster). Samples with clustered breakpoints within the same chromosome are classified as a single integration locus if the 5’ and 3’ most breakpoints are within 3 Mb of each other. (**b**), Schematic representation of clustered breakpoints at CESC integration loci across the human genome. Lines connecting to each chromosome represent different integration loci. Blue circles represent the indicated number of breakpoints per integration locus; orange boxed regions represent integration hotspots. See **Supplementary Figure S3** for the distribution of clustered breakpoints at integration loci in HNSCC tumors. (**c**), Scatter plot showing the frequency of single and clustered breakpoints per integration locus for CESC tumors grouped according to whether they overlap integration hotspots. The p-value is based on a non-parametric, unpaired t-test (two-tailed; **P<0.01).

Sites of recurrent HPV DNA integration in different tumor samples are termed integration hotspots. We defined integration hotspots (five or more sites located <5Mb apart) in our CESC dataset and compared them to previously defined hotspots from the literature ^10,15,28,36–40^. We identified a total of 37 hotspots in CESC tumors (**Supplementary Table S4**), which represented 313/584 (53.6%) integration loci from our CESC dataset (**Figure 2b-c**). Twenty-three hotspots overlapped previously defined sites of recurrent integration and 14 were novel hotspots, **Supplementary Figure S2** and **Supplementary Table S4-S5**. We were unable to define sites of recurrent integration in the HNSCC dataset because of the low number of integration loci in these tumors.

The distribution of clustered breakpoints at each integration locus and across each chromosome is shown in **Figure 2b** for CESC samples and **Supplementary Figure S3** for HNSCC samples. Most integration sites had 1-2 breakpoints in both the CESC (81.1%) and HNSCC (75.9%) datasets. Sites of recurrent integration are indicated by orange boxes for the CESC samples. The integration loci at these hotspots were more likely to contain clustered breakpoints compared to integration sites elsewhere in the genome for CESC tumors (**Figure 2c**; p=0.004). Higher numbers of clustered breakpoints at sites of recurrent integration suggests that these regions are susceptible to genomic instability.

### Most tumors with integrated HPV DNA have a single driver integration

Constitutive expression of the viral oncogenes from the integration locus is required for clonal selection and oncogenic progression. Transcriptionally silent HPV integration loci can be considered to be passenger sites ^35^. To identify driver versus passenger integrations, the transcriptional activity of each integration locus was determined for the subset of samples that had matched RNA sequencing data (CESC, n=144; HNSCC, n=35). The transcription status of each integration locus is indicated in **Supplementary Table S2-S3**. Integration loci in which no HPV mRNA was detected by RNA-seq analysis were classified as inactive or passenger loci. In CESC, all samples with a single integration locus (n=86) were transcriptionally active (**Figure 3a**). For samples with multiple integration loci (CESC, n=58; HNSCC, n=13), more than one transcriptionally active integration locus was observed in 35 (60.3%) CESC and four (30.8%) HNSCC tumors. Three HNSCC samples had no driver integrations; one sample (4.5%) had a single integration locus and two samples (15.4%) had multiple integration loci, **Figure 3b**. Overall, the majority of CESC and HNSCC tumors with integrated HPV genomes had only a single transcriptionally active integration locus. This implies that most tumors with integrated HPV DNA have a single driver integration. Viral oncogene transcription was analyzed at integration hotspots in CESC, which showed that sites of recurrent integration can have both driver and passenger integrations but are more likely to be transcriptionally active (**Figure 3c**).

**Figure 3.**
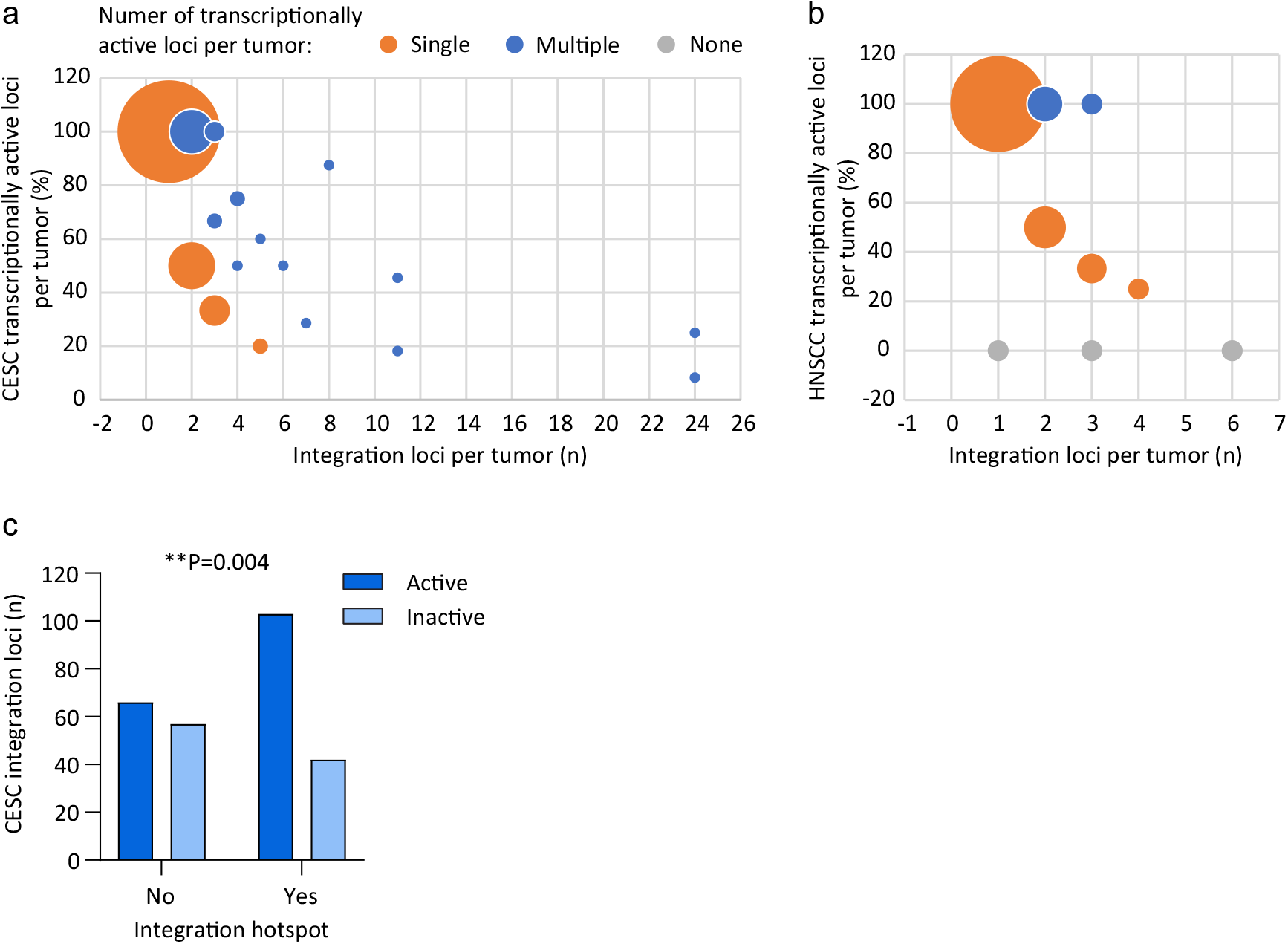
Transcription status of integrated viral genomes. (**a-b**), Bubble graph showing the percentage of transcriptionally active integration loci per tumor in CESC (**a**) and HNSCC (**b**) samples relative to the number of integration loci per tumor; 100% indicates that all integration loci are active in that tumor. Orange, blue and grey circles represent tumors with a single, multiple or no transcriptionally active loci, respectively. Circle size indicates the number of samples per grouping (for CESC, largest, n=86 and smallest, n=1; for HNSCC, largest, n=21 and smallest, n=1). Three HNSCC samples (*TCGA-CR-6482*, *TCGA-CN-5374* and *TCGA-CR-7404*) were reported as integration negative from RNA-seq ^30^ but had a single or multiple integration loci detected through WGS ^5^ and were therefore classified as transcriptionally inactive. (**c**), Bar chart showing the number of CESC integration loci that are transcriptionally active or inactive for viral oncogene expression at integration hotspots. Association between viral oncogene transcription and integration hotspots was based on a Fisher’s exact test (two-tailed; **P<0.01).

### Clustered integration breakpoints are associated with amplified regions of the host genome in CESC and HNSCC

HPV integration loci often have amplification and/or rearrangements of the flanking cellular sequences at the insertion sites ^6,7,9,15^. We determined the frequency with which integration loci in our datasets were associated with somatic copy number alterations for the subset of samples that had matched host genome copy number data (235 CESC and 22 HNSCC tumors; **Supplementary Table S6-S7**). In this subset, most integration breakpoints occurred within amplified regions of cellular DNA in both CESC and HNSCC relative to regions with a normal genomic copy number or adjacent to deletions (**Figure 4a-b**). In addition, the number of breakpoints per integration locus was significantly different at amplified regions of the host genome relative to loci with a normal genomic profile for both CESC and HNSCC samples; higher numbers of breakpoints per cluster occurred at amplified regions (**Figure 4a-b**). No significant difference was found in the number of clustered breakpoints per integration loci in regions with a normal genomic copy number relative to those with genomic copy number losses in CESC samples. We conclude that integration loci associated with flanking host DNA amplification were more likely to contain clustered breakpoints and to have higher numbers of breakpoints per integration locus.

**Figure 4.**
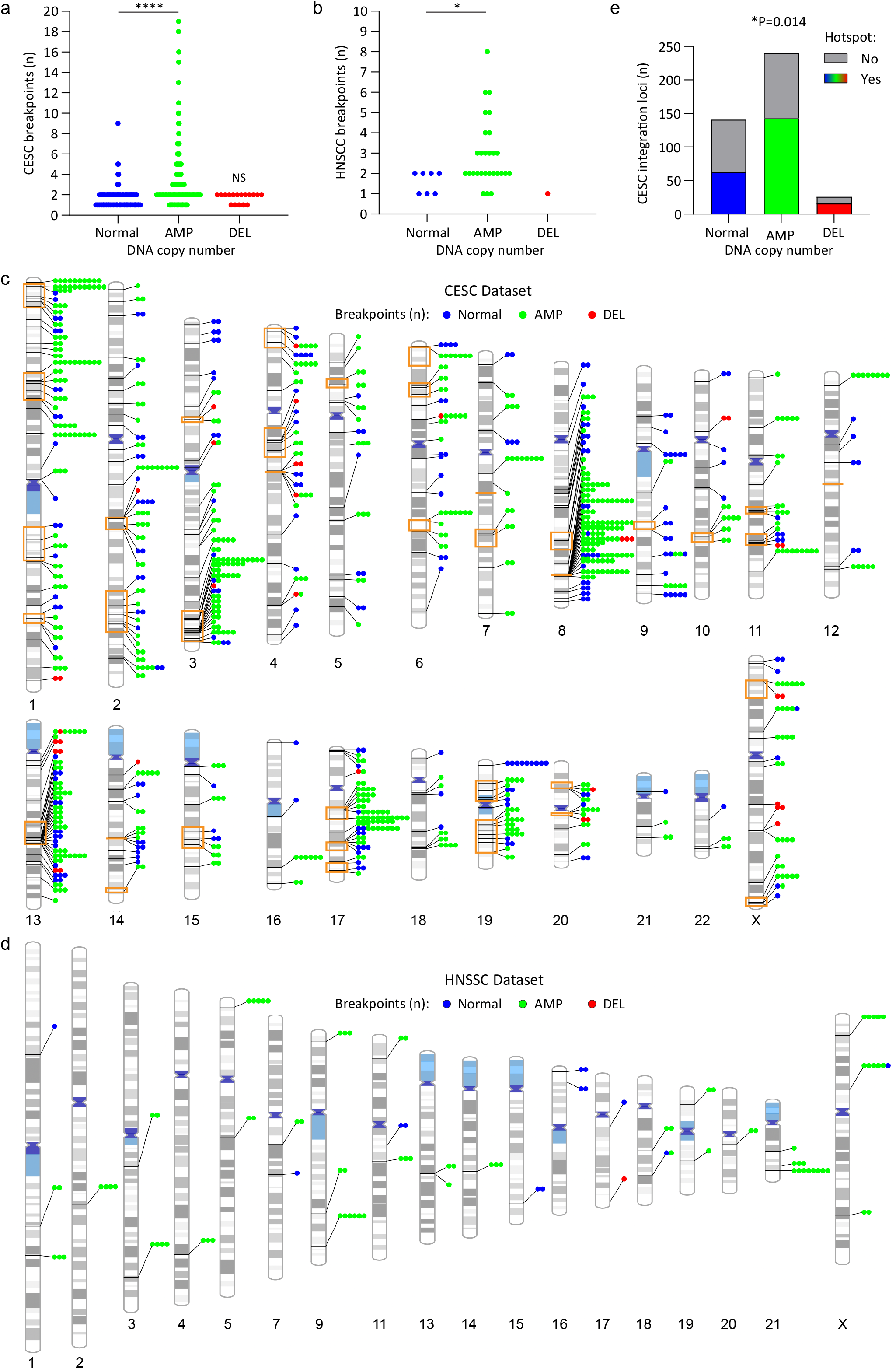
Clustered integration breakpoints are associated with amplified regions of the host genome in CESC and HNSCC. For the subset of CESC and HNSCC samples that had matched somatic copy number alteration data, HPV integration breakpoints were grouped according to the associated host DNA copy number status. Normal, AMP (amplification) and DEL (deletion) refers to the genomic profile of the host DNA at the integration locus. (**a-b**), Scatter plots showing the number of breakpoints per locus grouped according to the somatic copy number alteration status of the integration locus for CESC (**a**) and HNSCC (**b**) tumors. For CESC, the number of integration loci per grouping was Normal, n=140; AMP, n=240 and DEL, n=17. For HNSCC, the number of integration loci per grouping was Normal, n=7; AMP, n=28 and DEL, n=1. P-values are based on non-parametric, unpaired t-tests (two-tailed; *p<0.05, ****p<0.0001, NS=Non-significant). All statistical tests were performed relative to integration loci with a normal genomic profile. DNA amplification associated with integration loci ranged from 693 bp to 54.2 Mb (average, 1.6 Mb; median 47.4 Kb) in CESC and 6.5 Kb to 102.7 Mb (average, 3.3 Mb; median 43.1 Kb) in HNSCC. (**c-d**), Schematic representation of clustered breakpoints at integration loci that have associated host somatic copy number alterations. Lines connecting to each chromosome represent different integration loci for the CESC (**c**) and HNSCC (**d**) datasets. The number of circles represents the number of breakpoints per integration locus. Blue, green and red colored circles respectively represent integration sites that have a normal genomic profile or associated amplifications or deletions. Orange boxed regions represent integration hotspots. (**e**) Stacked bar chart showing the number of CESC integration loci that overlap integration hotspots grouped according to whether they have associated somatic copy number alterations. Association between somatic copy number alterations and integration hotspots was based on a chi-square test (*p<0.05).

Integration loci with associated focal amplifications were found on all chromosomes in CESC and on 15 chromosomes in HNSCC. The distribution of clustered integration breakpoints relative to host genome amplification and sites of recurrent integration in CESC is shown, **Figure 4c-d**. There was an enrichment of integration loci with associated host DNA amplifications at integration hotspots in CESC (**Figure 4e**) and higher numbers of clustered breakpoints were common at these regions (**Figure 4c**). Genomic instability in these regions most likely results in co-amplification of both viral and host DNA.

### Identification of common fragile sites in cervical cancer cell lines

Genomic instability occurs frequently at common fragile sites. We previously used FANCD2 ChIP-chip to map fragile sites in an aphidicolin-treated cervical carcinoma cell line (C33-A) ^26^. Here, we have extended the FANCD2 dataset using ChIP-seq analysis of both C33-A and HeLa cervical carcinoma cells (**Supplementary Table S8-S9**) and combined these results with those previously mapped in C33-A cells by ChIP-chip ^26^. There was good overlap between the FANCD2 peaks called across the three datasets (p<0.0001). In total we defined 513 FANCD2-enriched regions between the two cervical carcinoma cell lines, and they are listed in **Supplementary Table S10**.

We compared our cervical carcinoma cell line derived FANCD2 mapped fragile sites (genomic coverage 7.9%) to the 77 aphidicolin-induced common fragile sites (FRA regions) defined cytogenetically in lymphocytes and reported in the HGNC (HUGO Gene Nomenclature Committee) database (genomic coverage 48.3%), **Figure 5a**. A total of 115 (22.4%) FANCD2-enriched regions derived from C33-A and HeLa cells overlapped with 55.8% (n=43/77) FRA-regions, **Figure 5b**. FRA-regions that overlapped with FANCD2-associated fragile sites are listed in **Supplementary Table S11**. Permutation testing was used to determine the significance in overlap between our FANCD2-dataset and traditional FRA regions. The association between these genomic features did not reach significance (p=0.0634), which reflects differences in replication stress at these regions in different cell types.

**Figure 5.**
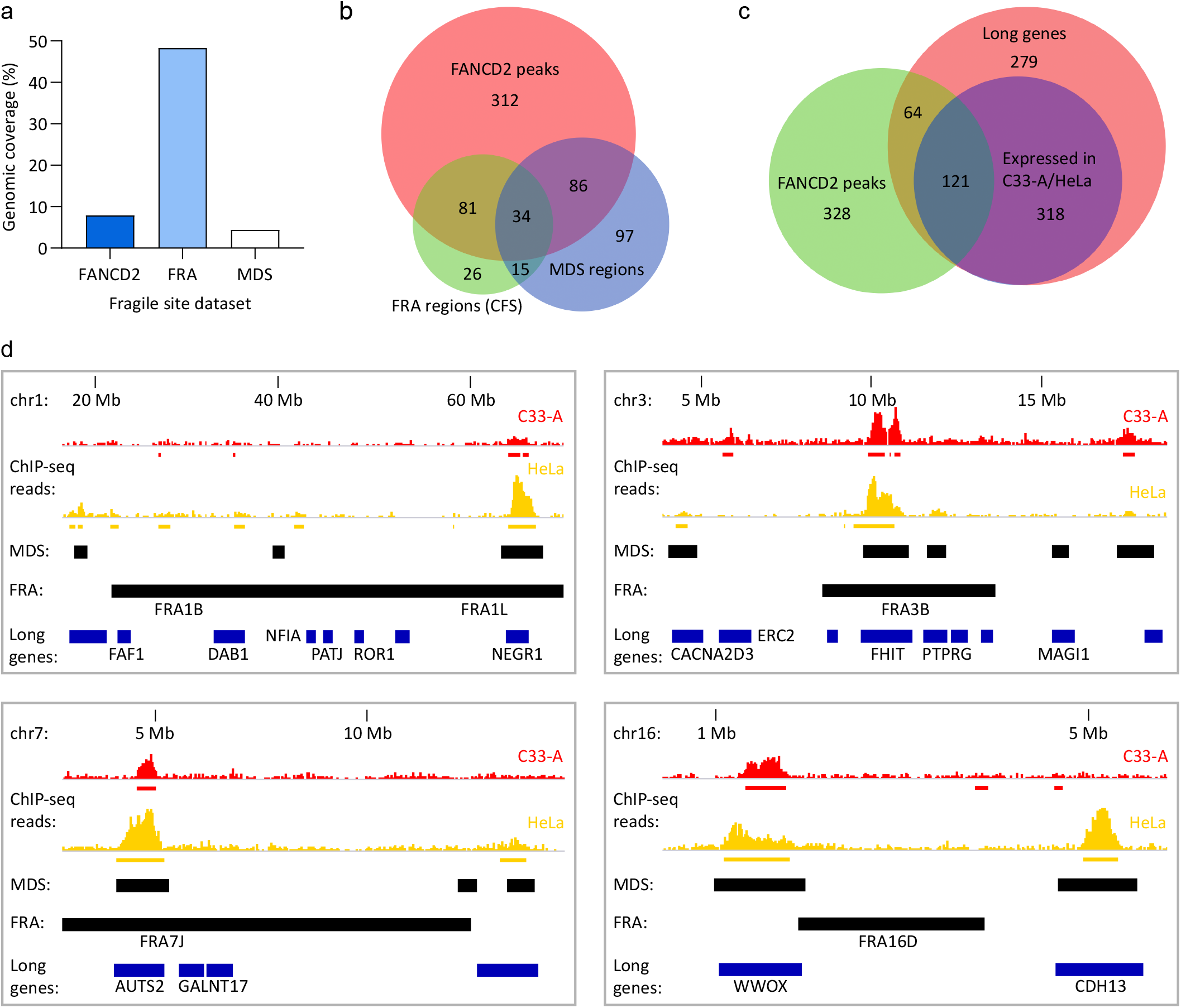
Cervical keratinocyte-specific fragile sites mapped by FANCD2 ChIP-seq. HPV-negative (C33-A) and HPV18-positive (HeLa) cervical carcinoma cells were treated for 24 hours with 0.2 µM aphidicolin and FANCD2-enriched regions were identified by ChIP-seq. All analyses were performed using the combined C33-A and HeLa FANCD2 dataset. (**a**), Bar graph showing the genomic coverage of FANCD2-enriched regions relative to common fragile sites (FRA regions) and mitotic DNA synthesis (MDS) regions reported in the literature ^41,42^. (**b**), Venn diagram showing the regions of overlap between fragile sites identified in the FANCD2, FRA and MDS datasets. (**c**), Venn diagram showing the overlap of FANCD2-associated fragile sites with protein-coding genes longer or equal to 0.3 Mb. Red circle represents all long genes; blue circle represents long genes that are expressed in C33-A and/or HeLa cells. (**d**), Alignment of FANCD2-enriched regions in C33-A (red) and HeLa (yellow) cells with associated genes (blue bars represent genes >0.3 Mb; genes expressed in C33-A and/or HeLa cells are indicated by gene name) and FRA and MDS regions (black bars). Red and yellow bars below the FANCD2 ChIP-seq signal tracks represent peaks mapped by SICER analysis in the corresponding cell lines.

Recent studies used high-resolution MiDA-seq (next-generation sequencing of EdU incorporation at sites of mitotic DNA synthesis) to map fragile sites in HeLa cells ^41,42^ and show that they colocalize with FRA-regions and FANCD2 foci in cells treated with aphidicolin. We compared the overlap between our dataset of FANCD2-enriched regions and the mitotic DNA synthesis regions profiled in HeLa cells (total genomic coverage of 4.4%, **Figure 5a**). A total of 120 (23.4%) FANCD2-enriched regions overlapped with mitotic DNA synthesis regions, which represented 48.3% (n=112/232) of mitotic DNA synthesis regions, **Figure 5b**. Permutation testing was used to determine the significance in overlap between FANCD2-enriched regions and mitotic DNA synthesis regions (p<0.0001). Mitotic DNA synthesis regions that overlapped with FANCD2-associated fragile sites are indicated in **Supplementary Table S12**. Collectively, these data show good correlation between our FANCD2-associated fragile sites and regions of the genome susceptible to genetic instability.

The instability of fragile sites is often due to transcription-replication conflicts that frequently occur at long genes ^43^. We and others have previously shown that FANCD2-enriched regions overlap with transcriptionally active long genes in C33-A ^26^ and U2OS cells ^44^. Here we extended that association to include genes that are >0.3 Mb in length ^45^. A total of 184/513 (35.9%) FANCD2-enriched regions overlapped with protein-coding genes that were ≥0.3 Mb, which corresponded to 185/782 (23.7%) long genes. A Fisher’s exact test was used to determine the significance in overlap between FANCD2-enriched regions and long genes (two-tailed, p=4.22E-13). Of the long genes that overlapped with FANCD2-enriched regions, 121/185 (65.4%) were expressed in C33-A and/or HeLa cells, **Figure 5c**, further validating our FANCD2 peaks as sites of genetic instability in cervical carcinoma cells. **Supplementary Table S13** lists the long genes used in this analysis and their association with common fragile sites and expression in C33-A and HeLa cells from RNA-seq (^26^ and Expression Atlas, https://www.ebi.ac.uk/gxa/home). Example alignments of our FANCD2-associated fragile sites relative to common FRA-regions, mitotic DNA synthesis regions and long genes are shown in **Figure 5d**.

### Integration breakpoints are enriched at FANCD2-associated fragile sites

To assess the association of our cervical carcinoma-specific fragile site dataset with HPV integration sites, we calculated the frequency with which an integration breakpoint occurred within 50 Kb ^9^ of the C33-A and HeLa FANCD2-enriched regions (**Supplementary Table S10**). Approximately 18% integration breakpoints were associated with FANCD2-enriched regions in both the CESC and HNSCC datasets. The data was permutated 10,000 times to create an expected distribution of the overlap between integration breakpoints and FANCD2-enriched regions. This showed that the FANCD2-associated fragile sites were significantly enriched for both CESC and HNSCC HPV integration sites when each breakpoint was analyzed independently from each other, **Table 1**.

**Table 1.**
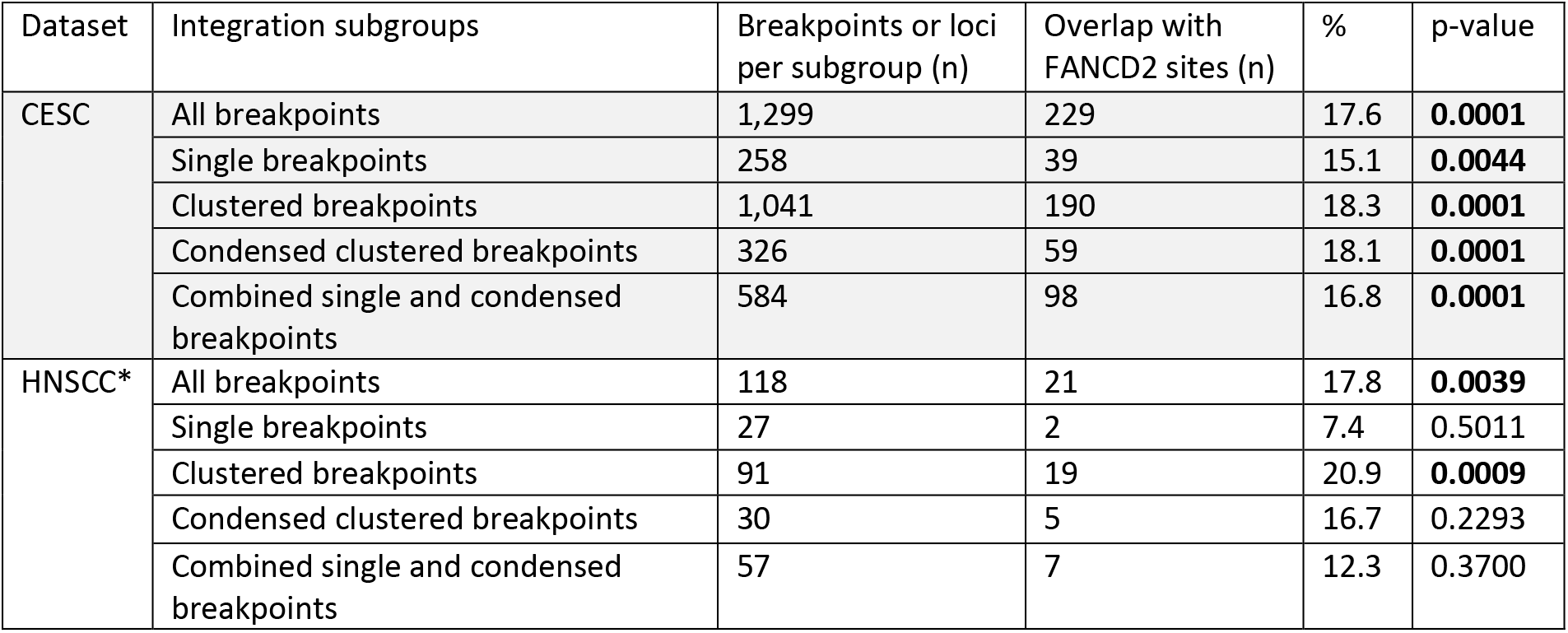
Overlap of integration breakpoints with FANCD2-assocated fragile sites. CESC and HNSCC integration breakpoints (−/+ 50 Kb flank regions) were grouped by the number of breakpoints per integration locus; single indicates one breakpoint and clustered indicates two or more breakpoints. Integration breakpoints from each subgroup were intersected with FANCD2-enriched regions and the frequency of overlap calculated. For the *All breakpoints*, *Single breakpoints (not clustered)* and *Breakpoints located within a cluster subgroups*, each breakpoint was tested independently for its overlap with FANCD2-enriched regions, regardless of whether it was part of a cluster or not. For the *A single cluster of breakpoints (using 5’-3’ endpoints)* subgroup, the region spanning the most 5’ and 3’ breakpoints of an integration locus was used to test for the overlap with FANCD2-enriched regions. For the *Single and clustered 5’-3’ breakpoints* subgroup, the *Single breakpoints (not clustered)* and *A single cluster of breakpoints (using 5’-3’ endpoints)* subgroups were combined for overlap analysis. The data was permutated 10,000 times to create an expected distribution of overlap. *A single integration breakpoint on chromosome Y was excluded from this analysis. Bold font indicates significant p-values.

| Dataset | Integration subgroups | Breakpoints or loci per subgroup (n) | Overlap with FANCD2 sites (n) | % | p-value |
| --- | --- | --- | --- | --- | --- |
| CESC | All breakpoints | 1,299 | 229 | 17.6 | **0.0001** |
|  | Single breakpoints | 258 | 39 | 15.1 | **0.0044** |
|  | Clustered breakpoints | 1,041 | 190 | 18.3 | **0.0001** |
|  | Condensed clustered breakpoints | 326 | 59 | 18.1 | **0.0001** |
|  | Combined single and condensed breakpoints | 584 | 98 | 16.8 | **0.0001** |
| HNSCC* | All breakpoints | 118 | 21 | 17.8 | **0.0039** |
|  | Single breakpoints | 27 | 2 | 7.4 | 0.5011 |
|  | Clustered breakpoints | 91 | 19 | 20.9 | **0.0009** |
|  | Condensed clustered breakpoints | 30 | 5 | 16.7 | 0.2293 |
|  | Combined single and condensed breakpoints | 57 | 7 | 12.3 | 0.3700 |

To remove bias resulting from overrepresentation of integration loci containing clusters of breakpoints, the integration loci were simplified into defined subsets for significance testing. Integration loci contain either a single breakpoint or a cluster of breakpoints (**Figure 2a**). In the latter group, the clustered breakpoints were condensed into a single site represented by the most 5’ and 3’ breakpoints. The final category combined the single and condensed categories, in which each integration locus was represented just once. The integration loci in each category were tested independently for their association with FANCD2-enriched regions. For the CESC dataset, sites containing single or clustered breakpoints were significantly associated with FANCD2-enriched regions, **Table 1**. In contrast, only sites with clustered breakpoints reached significance for the HNSCC dataset (**Table 1**).

### Generation of a cervical keratinocyte enhancer dataset using Brd4 and H3K27ac ChIP-seq

It has been noted previously that HPV integration occurs frequently at transcriptionally active regions ^15,23^, and we have demonstrated that HPV integration can capture and amplify cellular enhancers to drive viral oncogene expression ^25^. Brd4 is a marker of cell lineage-specific enhancers ^46–48^ and HPV E2 tethering sites ^26,49^. Moreover, we, and others, have shown previously that Brd4 and the HPV E2 replication protein bind to transcriptionally active chromatin within the host genome ^49,50^ that overlap many FANCD2-associated fragile sites ^26^. Viral replication factories form adjacent to these sites and we have proposed that tethering of the viral genome to these unstable sites would increase the chances of integration at these regions.

To further examine the association of cellular enhancers with HPV integration in CESC and HNSCC, we generated an “in-house” enhancer dataset using Brd4 and H3K27ac ChIP-seq in four different subclones of W12 cervical keratinocytes. We defined 6,935 enhancer consensus peaks in the four W12 subclones (**Supplementary Table S14**). The resulting H3K27ac and Brd4 ChIP-seq signals were compared to enhancers in the NHEK (Normal Human Epidermal Keratinocytes) ENCODE dataset ^51^, which showed that 83.5% (p<0.0001) W12 Brd4/H3K27ac enriched peaks overlapped ENCODE defined enhancers (**Supplementary Figure S4**). Approximately 40% integration breakpoints and loci from the CESC and HNSC datasets were significantly associated with our cervical keratinocyte enhancer dataset, **Table 2**.

**Table 2.** Overlap of integration breakpoints with keratinocyte-specific enhancers. CESC and HNSCC integration breakpoints (−/+ 50 Kb flank regions) were grouped by the number of breakpoints per integration locus; single indicates one breakpoint and clustered indicates two or more breakpoints. Integration breakpoints from each subgroup were intersected with cellular enhancers and the frequency of overlap calculated. For the *All breakpoints*, *Single breakpoints (not clustered)* and *Breakpoints located within a cluster* subgroups, each breakpoint was tested independently for its overlap with enhancer regions, regardless of whether it was part of a cluster or not. For the *A single cluster of breakpoints (using 5’-3’ endpoints)* subgroup, the region spanning the most 5’ and 3’ breakpoints of an integration locus was used to test for the overlap with enhancer regions. For the *Single and clustered 5’-3’ breakpoints* subgroup, the *Single breakpoints (not clustered)* and *A single cluster of breakpoints (using 5’-3’ endpoints)* subgroups were combined for overlap analysis. The data was permutated 10,000 times to create an expected distribution of overlap. *A single integration breakpoint on chromosome Y was excluded from this analysis. Bold font indicates significant p-values.

| Dataset | Integration subgroups | Breakpoints or loci per subgroup (n) | Overlap with Enhancers (n) | % | p-value |
| --- | --- | --- | --- | --- | --- |
| CESC | All breakpoints | 1,299 | 495 | 38.1 | **0.0001** |
|  | Single breakpoints | 258 | 67 | 26.0 | **0.0001** |
|  | Clustered breakpoints | 1,041 | 428 | 41.1 | **0.0001** |
|  | Condensed clustered breakpoints | 326 | 162 | 49.7 | **0.0001** |
|  | Combined single and condensed breakpoints | 584 | 229 | 39.2 | **0.0001** |
| HNSCC* | All breakpoints | 118 | 43 | 36.4 | **0.0001** |
|  | Single breakpoints | 27 | 9 | 33.3 | **0.0052** |
|  | Clustered breakpoints | 91 | 34 | 37.4 | **0.0001** |
|  | Condensed clustered breakpoints | 30 | 13 | 43.3 | **0.0011** |
|  | Combined single and condensed breakpoints | 57 | 22 | 38.6 | **0.0003** |

### Integration hotspots are associated with gene loci related to cell development and identity

Enhancers that were associated with integration loci were analyzed using the Genomic Regions Enrichment of Annotations Tool (GREAT) ^52^ to identify common functional significance based on proximity testing. This identified 52 putative enhancer target genes for the CESC dataset, representing 28 gene loci of which 60.7% overlapped integration hotspots (**Figure 6a**). Twelve of the gene loci are previously defined sites of recurrent integration and include the KLF5, KLF12, MYC, TP63, RAD51B and HMGA2 genes, and five of the gene loci are novel integration hotspots and include the CAMK1G, FOXQ1, EXOC2, GRHL2, ID1, COX4I2, HM13 and NFIA genes (**Figure 6a**). For HNSCC, 19 target genes were identified through GREAT gene ontology analysis, representing eight gene loci of which 50% overlapped integration hotspots profiled in CESC, **Supplementary Figure S4**. Six target genes (KLF5, KLF12, FAM84B, POU5F1B, TUBD1 and VMP1) were common across CESC and HNSCC. Gene ontology analysis of our enhancer regions identified epithelium development, epithelial cell differentiation, and negative regulation of keratinocyte differentiation to be significantly enriched biological processes associated with CESC integration loci (**Supplementary Table S15**). Ectoderm development and differentiation, epidermal and epithelial differentiation, tongue morphogenesis and negative regulation of spreading epidermal cells during wound healing were biological processes significantly associated with enhancer enrichment at HNSCC integration loci, **Supplementary Table S15**. This indicates that sites of recurrent HPV integration are often associated with cellular pathways relevant to host cell development and differentiation.

**Figure 6.**
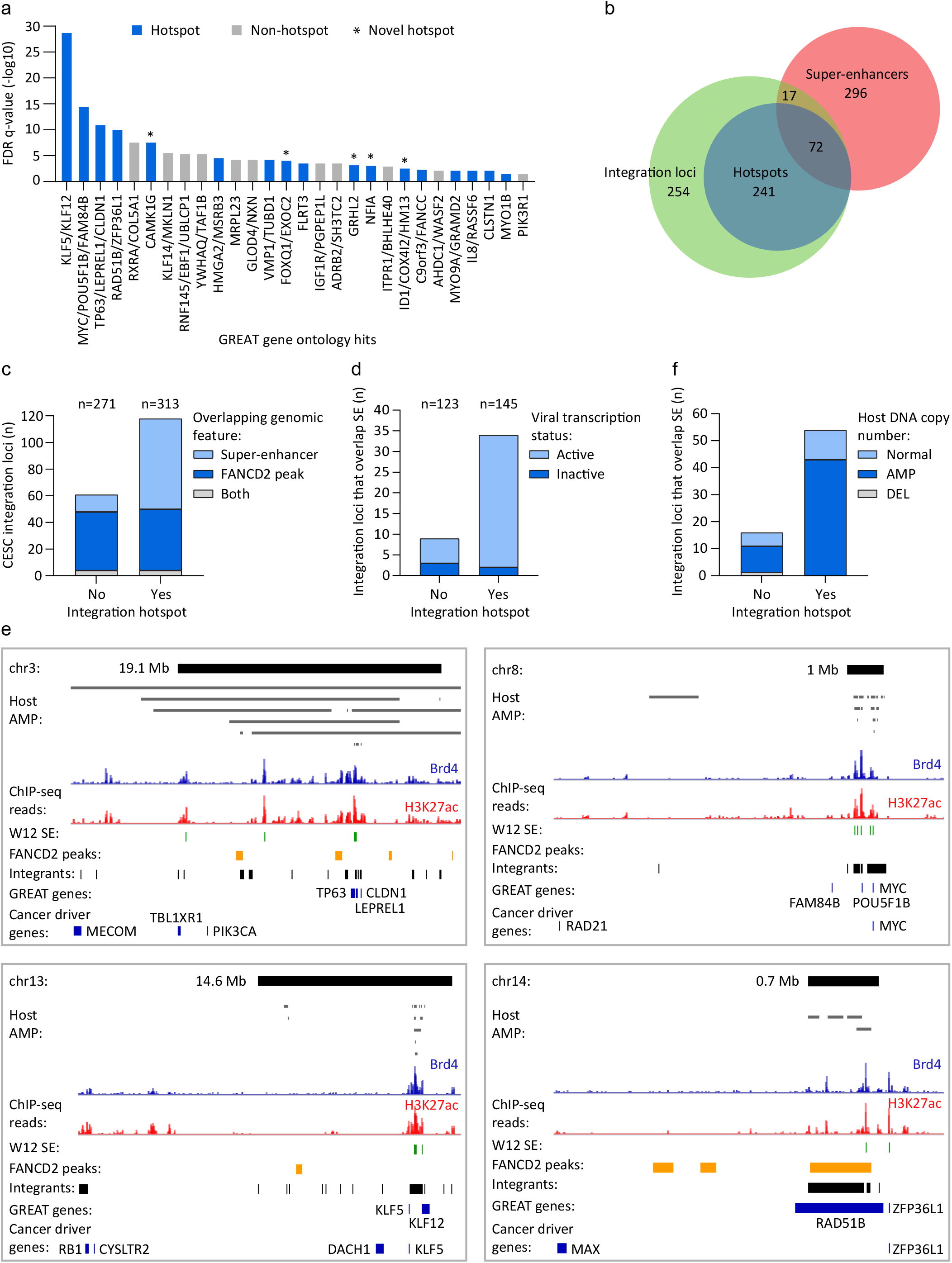
Integration hotspots are associated with cellular super-enhancers. H3K27ac and Brd4 enriched regions were profiled in HPV16-positive cervical derived W12 keratinocyte subclones by ChIP-seq. Enhancer regions were defined as peaks that overlapped in both H3K27ac and Brd4 datasets, and that were identified across the four W12 subclones. (**a**), GREAT (Genomic Regions Enrichment of Annotations Tool) gene ontology analysis was performed using W12 enhancers that overlapped CESC integration breakpoints (−/+ 50 Kb flanks) as input, and compared against all W12 enhancers, to identify putative target genes associated with these cis-regulatory regions based on enhancer frequency. Bars represent putative target genes plotted against their FDR (false discovery rate) adjusted p-values (q-value). Blue and grey bars represent genes that overlap integration hotspots and sites of non-recurrent integration, respectively. Enriched target genes within the same genomic locus were grouped (e.g. KLF5 and KLF12) and plotted using the most significant q-value. (**b**), Venn diagram showing the regions of overlap between integration loci, integration hotspots and super-enhancers mapped in W12 subclones. (**c**), Bar chart showing the number of CESC integration loci that were grouped according to whether or not they are integration hotspots and plotted based on their overlap with super-enhancers, FANCD2-associated fragile sites or both genomic features. Numbers above the graph indicate the total number of integration loci within each grouping. (**d**), Bar chart showing the number of CESC integration loci that are associated with super-enhancers (SE) that were grouped according to whether they are integration hotspots and plotted based on their viral transcription status. Numbers above the graph indicate the total number of integration loci that have associated viral transcription data within each grouping. (**e**), Alignment of Brd4 (blue) and H3K27ac (red) ChIP-seq signals mapped in W12 cervical keratinocytes at integration hotspots (top black bars; size indicated in Mb) in cervical carcinomas. Grey bars represent amplified (AMP) host DNA in different CESC tumors from The Cancer Genome Atlas. Green, yellow, and black bars below the ChIP-seq signal tracks represent super-enhancers (SE) mapped in W12 subclones, FANCD2-associated fragile sites mapped in C33-A and HeLa cells and CESC integration loci, respectively. Genes identified from GREAT gene ontology analysis ^52^ and cancer driver genes ^55^ are indicated by blue bars. Each integration hotspot is characterized in **Supplementary Figure S5** and **Table S4**. (**f**), Bar chart showing the number of CESC integration loci that are associated with super-enhancers (SE) that were grouped according to whether they are integration hotspots and plotted based on their host somatic copy number alternation status. The number of integration loci per grouping was normal, n=140; amplification (AMP), n=240 and deletion (DEL), n=17.

### Keratinocyte-specific super-enhancers are enriched at integration loci

Large enhancer clusters were characteristic of the integration targets identified through GREAT analysis. We therefore defined super-enhancers in our Brd4-defined W12 enhancer dataset based on relative peak height of the H3K27ac and Brd4 ChIP-seq datasets using the Rank Ordering of Super-Enhancers (ROSE) tool ^53,54^. We defined 338 super-enhancers in W12 cervical keratinocytes (**Supplementary Table S16**). Intersect analysis showed that 89/584 (15.2%) CESC integration loci overlapped with super-enhancers profiled in W12 cells, and of these loci 72 (80.9%) were classified as sites of recurrent integration (**Figure 6b**). A total of 25/37 (67.6%) integration hotspots contained super-enhancers and permutation testing showed that both CESC integration loci and sites of recurrent integration were significantly associated with these regulatory domains (p<0.0001). For HNSCC, 8/57 (14%) integration loci were associated with W12 super-enhancers, and 62.5% of these loci overlapped integration hotspots profiled in CESC, including the KLF5/KLF12, MYC, ERBB2 and VMP1 gene loci. Permutation testing showed that HNSCC integration loci were significantly associated with super-enhancers profiled in W12 cells (p<0.0001). Thus, keratinocyte-specific super-enhancers are enriched at integration loci in HPV-associated tumors and are frequently found at integration hotspots.

The association of CESC integration loci with super-enhancers and FANCD2-enriched fragile sites at integration hotspots was also addressed, **Figure 6c**. This showed that the frequency of FANCD2 enrichment at integration loci was comparable for sites of recurrent (50/313 loci; 16.0%) and non-recurrent integration (48/271 loci; 17.7%) in CESC, whereas the association of super-enhancers was augmented at integration loci that occurred within hotspots (72/313 loci; 23.0%) relative to non-recurrent sites of integration (17/271 loci; 6.3%). Most integration loci that overlapped super-enhancers were active for viral oncogene expression (driver integrations) and were more frequently observed at integration hotspots, **Figure 6d**. Furthermore, several cancer driver genes, including ASXL1, CACNA1A, IRF6, KANSL1, KLF5, KRT222, MYC, PPM1D, PTCH1 and PTPDC1 ^55^ were located within 1 Mb of super-enhancers that overlapped with integration hotspots, **Supplementary Figure S5** and **Table S17**. Alignment of FANCD2-associated fragile sites, super-enhancers and associated target genes at integration hotspots are shown in **Figure 6e** and **Supplementary Figure S4-S5**. Collectively, these data show that transcriptionally active chromatin and/or regions of genetic instability are common features of HPV integration sites. Moreover, integration hotspots are commonly associated with super-enhancers, several of which regulate cancer driver and/or cell-identity genes.

### Super-enhancers are frequently amplified at integration hotspots in CESC

The association of super-enhancers at integration hotspots was compared with the host somatic copy number alteration in CESC samples. Super-enhancers were more frequently observed at those CESC integration loci with associated host DNA amplifications (53/240; 22.1%) relative to those that had either a normal genomic profile (16/140; 11.4%) or deletions within the host DNA flanking sequences (1/17; 5.9%), **Figure 6f**. Of the amplified CESC integrations that had associated super-enhancers, 43 (81.1%) were sites of recurrent integration and represented 43/143 (30.1%) hotspot and 10/97 (10.3%) non-hotspot loci with associated host genome amplifications, **Figure 6f**. Super-enhancer overlap was also more frequently observed at integration hotspots (11/62; 17.7%) than non-hotspots (5/78; 6.4%) for loci with a normal genomic profile, representing 68.8% of loci that overlapped super-enhancers for this subgroup. However, for integration loci that had associated host deletions, no super-enhancers were observed at sites of recurrent integration (**Figure 6f**). This data shows that amplification of super-enhancers is frequently observed at integration loci in CESC, particularly at sites of recurrent integration.

## DISCUSSION

Many studies have documented the “landscape” of HPV integration sites with respect to traditional common fragile sites, host genome amplification, and transcription and related regulatory elements ^9–13,15,23^. Here, we combined and curated DNA and RNA sequencing datasets from HPV-positive CESC and HNSCC tumors and compared them with novel “in-house” datasets of common fragile sites defined by FANCD2 ChIP-seq, and enhancers and super-enhancers defined by Brd4 and H3K27ac ChIP-seq in cervical carcinoma derived cells. We show that viral integration sites in CESC are enriched at FANCD2-associated fragile sites in cervical cells. We also show that cervical cell enhancers are over-represented at HPV integration sites and that HPV integration is often associated with super-enhancers, particularly at integration hotspots enriched for cell-identity genes. Furthermore, we show that the flanking host DNA that is enriched for enhancers and super-enhancers is frequently amplified in CESC tumors.

HPV genomes replicate as extrachromosomal nuclear minichromosomes at every stage of the infectious cycle. The virus relies on the host replication and transcriptional machinery and it is thought that the HPV genome localizes to regions of the nucleus that facilitate these processes ^56^. At different stages of infection, the viral DNA associates with nuclear ND10 bodies, and interphase and mitotic host chromatin, and highjacks the DNA damage repair processes to amplify viral DNA ^57,58^. Concomitantly, the viral E6 and E7 proteins induce cell proliferation and replication stress, abrogate cell cycle checkpoints, and inhibit the innate immune response ^59^. E6 and E7 proteins also modify the epigenetic landscape of the host genome by changing the levels of different histone-modifying enzymes ^60^. Together, these activities could promote the accidental integration of viral DNA that is closely associated with host chromatin.

HPV genomes replicate using two different modes: in maintenance replication the genomes replicate bidirectionally at low copy number, but this switches to a unidirectional recombination-directed mechanism in the amplification stage ^61,62^. The formation of tandem repeats at integration sites could be related to these processes and over-replication of viral and host sequences could result from repeated initiation of replication at the viral replication origin, especially if the HPV E1 and E2 proteins are expressed ^21,63^. In fact, unscheduled firing of replication origins and increased replication fork stalling has been shown to occur in both viral and host sequences at HPV integration sites in the MYC locus ^20^, which is frequently amplified in HPV-associated cancers. Tandem repeating units of co-amplified viral and cellular DNA could result from this endoreduplication, replication fork arrest, and homologous recombination. Highly rearranged integrations are also consistent with the breakage-fusion-bridge–type model of genome amplification ^19^. At fragile sites, perturbed replication dynamics could also generate focal amplifications and/or rearrangements of viral-host sequences.

In this study, we generated a dataset of aphidicolin-induced common fragile sites in two cervical carcinoma cell lines, C33-A and HeLa, and found a significant association between these sites and integration breakpoints in CESC, particularly at those loci with clustered breakpoints. Common fragile sites are susceptible to somatic copy number alterations ^64,65^ likely due to replication stress that arises from perturbed replication dynamics in conflict with transcription of long genes ^66^. Accordingly, our C33-A and HeLa common fragile sites were overrepresented at long genes expressed in these cells. We did not observe an enrichment of HPV integration sites in HNSCC samples with our FANCD2-associated common fragile sites, although an association was previously noted between traditional FRA-regions and integration sites in oropharyngeal squamous cell carcinomas ^10^. This difference could reflect the larger genomic coverage of FRA-regions used in the previous study, or the limited number of samples in our HNSCC dataset. Moreover, common fragile sites are likely distinct in cervical and oropharyngeal derived keratinocytes, or alternatively these findings could reflect differences in the biology of HPV infection and mechanisms of oncogenic progression in the different tissue types.

HPV integration often occurs in transcriptionally active chromatin within the host genome ^15,23,67^ and we previously described an example of enhancer-hijacking and co-amplification of cellular and viral regulatory sequences at an HPV integration site in cervical lesion derived cells ^25^. The association of viral integration breakpoints with putative enhancer regions in HPV-associated cancers has been reported ^15^, but the enhancer regions used were based on ENCODE histone modifications and therefore did not reflect the specific enhancer profiles of HPV-positive cervical cells. Here, we defined keratinocyte enhancers in HPV16-positive W12 cervical keratinocytes by H3K27ac and Brd4 enrichment and show that these specific enhancers are significantly overrepresented at HPV integration loci. In some cases, these loci were associated with focal amplification of host DNA, providing evidence for potential enhancer-capture. A recent study showed enrichment of active histone marks at HPV integration loci in cervical tumors, which correlated with upregulation of local gene expression, and increased gene expression levels at loci with increased breakpoints ^68^. Kamal et al. also found increased local host gene expression at loci with multiple junction copies (analogous to clustered breakpoints) ^36^. Furthermore, integration loci with associated somatic copy number alterations have also been shown to have increased gene expression ^29^. These observations could represent enhancer-hijacking. Integration hotspots often contain genes that drive cancer ^55^ and accidental integration at these regions could results in clonal expansion and selection due to perturbation of these oncogenic pathways. Thus, enhancer-hijacking can drive expression of both viral and cellular genes.

Super-enhancers are large clusters of enhancers, rich in Brd4 binding and H3K27ac modification, that often control cell identity genes and are coopted in tumorigenesis ^53,54^. We defined super-enhancers in our W12 cervical cell line datasets and showed that they were strongly associated with integration hotspots, including the MYC, KLF5/KLF12 and ERBB2 gene loci, which are important regulators of cell-cycle, proliferation and apoptosis ^69–71^; TP63, which is a master regulator of epidermal keratinocyte proliferation and differentiation ^72^; and RAD51B, which is a key regulator of homologous recombination repair ^73^. We propose that, during persistent infection, extrachromosomal HPV genomes specifically localize at key transcriptional regulatory hubs within the host genome, several of which are important for keratinocyte biology.

The cellular Brd4 protein is involved in many of the cellular and viral processes described in this study. Brd4 is a chromatin scaffold protein that modulates transcriptional initiation and elongation and is a major component of super-enhancers ^74^. Brd4 is also important at multiple stages of the HPV infectious cycle ^75^, binds to common fragile sites in C33-A cells ^26^, and is important for tethering HPV genomes to mitotic chromatin ^76,77^. Brd4 is enriched at the HPV16 integration site/super-enhancer in W12 cells and inhibition of Brd4 binding reduces E6/E7 transcription and cell growth ^24^. Therefore, Brd4 is an example of a factor that is crucial for key cell and viral chromatin-related processes, and the juxtaposition of these processes could promote integration of viral DNA and oncogenic progression.

In conclusion, many factors contribute to the integration of a viral genome that eventually drives oncogenesis. Cancer genomes often contain multiple HPV integration sites but usually only one is transcriptionally active (this study, ^29,35^). In CESC-derived cells, expression of the viral E6 and E7 oncoproteins is necessary for cell proliferation, survival, and maintenance of the tumor phenotype ^78^. Therefore, integration is common, but requires the right genomic location for constitutive viral oncogene expression. For example, the viral oncogenes are usually expressed from a viral-host fusion transcript that requires a splice acceptor and polyadenylation signal in the flanking host DNA ^79,80^. Therefore, HPV integrants require a combination of events and processes that are dependent on the genetic and/or epigenetic landscape of the flanking host chromatin to drive oncogenesis.

## METHODS

### HPV integration datasets

A systematic literature review identified genomic datasets from HPV-positive CESC and HNSCC that contained information on HPV type and integration breakpoints within the host and viral genomes, which were identified by sequencing (**Supplementary Table S1**) ^5,6,8,9,28–35^. The integration breakpoints used in this study were originally identified from both RNA and DNA sequencing methods, including APOT (amplification of papillomavirus oncogene transcripts), DIPS (detection of integrated papillomavirus sequences) and next-generation sequencing technologies. The use of hybrid-capture technologies for detection of viral integration sites has been reported to give high rates of false positives ^33,34^, and so insertion breakpoints identified by this method were only included if they were validated by other means, such as Sanger sequencing. The methodology used to identify each integration breakpoint is referenced in **Supplementary Table S2 and S3.** RNA-based sequencing methods give an approximation of the insertion site based on the closest splice acceptor site within the host genome; therefore, integration breakpoints identified by RNA-seq and/or APOT were only used to determine the viral transcription status of an integration site for samples with matched DNA sequencing data. For TCGA CESC and HNSCC samples, unmapped reads were extracted from RNA-seq, whole genome sequencing (WGS) and whole exome sequencing (WXS) BAM files (https://portal.gdc.cancer.gov/legacy-archive; accessed 01/01/2013) and pre-processed with prinseq-lite.pl version 0.20.2 to remove low quality reads ^81^. Pre-processed reads were mapped with Bowtie 2 using the *very-sensitive* preset option against the Viral Refseq database ^82,83^. All unmapped reads were subjected to BLASTN with default parameters against the Viral Refseq database ^84^. All aligned reads were then subjected to BLASTN against the human hg19 reference assembly. Bowtie 2 and BLASTN reports were passed into SummonChimera using a 1,000 bp deletion size for integration detection ^85^. The SummonChimera reports were manually parsed to remove chimeric junctions with lower than 20 read coverage, chimeric junctions with no cross-analysis verification, and ambiguously reported integration predictions. Finally, a unique ID was provided to all uniquely detected chimeric junctions. For analysis of the association of integration breakpoints with different genomic features of interest, only integration breakpoints identified by DNA sequencing methods were used. The characteristics of samples included in this study, categorized by histology type, HPV type, tumor location and sequencing methods are summarized in **Supplementary Figure S1**. CESC and HNSCC integration breakpoints included in this study are listed in **Supplementary Table S2-S3**.

### Integration hotspot dataset

Integration loci from CESC tumors that were within 5 Mb of each other were collapsed into a single genomic interval to define integration hotspot boundaries. Exceptions to this size cut-off for collapsing adjacent integration loci were permitted to reflect previously defined hotspots from the literature. Five or more integration loci per hotspot (or three or more integration loci for sites that overlapped previously defined hotspots from the literature) were used to define sites of recurrent integration. Integration hotspots defined from our CESC dataset and previously in the literature are listed in **Supplementary Tables S4** and **S5**, respectively.

### Somatic copy number alteration datasets

The amplification status of the cellular sequences flanking integration breakpoints had been assessed in a subset of samples by comparative genomic hybridization (CGH) or SNP array datasets. For CGH array data, we defined two-fold or more focal amplification or deletion of the host genome at an integration locus as having an associated somatic copy number alterations ^8,28^. Matched SNP6 copy number segment data for TCGA CESC and HNSCC tumor samples were downloaded from the Broad Institute on 04/07/2020, http://firebrowse.org/ (genome_wide_snp_6-segmented_scna_minus_germline_cnv_hg19) ^86,87^. Positive and negative mean segment values above 0.3 and below −0.3 represented copy number gains and losses, respectively, and mean segment values between 0.3 and −0.3 were considered noise ^86,87^. Deletions that occurred within 50 Kb of an integration breakpoint were included in this analysis. Deletions that directly overlapped an integration breakpoint were excluded; these likely reflected the alternative chromosome as sequencing of the chimeric viral-host junction was available for the associated HPV insertion sites. Integration loci defined as having associated somatic copy number alterations in CESC and HNSCC are detailed in **Supplementary Table S6** and **S7**, respectively.

### Cell culture

Subclones (20831, 20861, 20862, 20863) derived from HPV16-positive W12 cervical keratinocytes ^79,88^ were maintained in F-medium (3:1 [vol/vol] F-12–Dulbecco’s modified Eagle’s medium, 5% fetal bovine serum, 0.4 μg/ml hydrocortisone, 5 μg/ml insulin, 8.4 ng/ml cholera toxin, 10 ng/ml epidermal growth factor, 24 μg/ml adenine, 100 U/ml penicillin and 100 μg/ml streptomycin). All cells were grown in the presence of irradiated 3T3-J2 feeder cells. C33-A and HeLa cervical carcinoma derived cell lines were maintained in Dulbecco’s modified Eagle’s medium, supplemented with 10% fetal bovine serum, 100 U/ml penicillin and 100 μg/ml streptomycin. To induce replication stress, C33-A and HeLa cells were treated for 24 hours with 0.2 µM aphidicolin (Sigma A0781) prior to harvesting for FANCD2 ChIP-seq experiments, described below.

### ChIP-seq: FANCD2

Aphidicolin treated C33-A and HeLa cells were processed for chromatin immunoprecipitation (ChIP) as previously described ^24^. Briefly, cells were crossed-linked with 1% formaldehyde and chromatin was isolated and sheared to 100–500 bp DNA fragments using a Bioruptor sonicator (Diagonode) on high power settings. Chromatin samples (25 μg per ChIP) were incubated overnight at 4°C with an antibody against FANCD2 (Bethyl, A302-174A, 2.5 μg). Rabbit IgG (Jackson ImmunoRes, 011-000-003) was used to determine non-specific binding to control regions (though not sequenced). Chromatin immunocomplexes were precipitated for 1 hour at 4°C with blocked Dynabeads Protein G (Invitrogen), subjected to multiple wash steps and the chromatin eluted in elution buffer (50 mM Tris-HCl [pH 8.0], 10 mM EDTA [pH 8.0], 1% SDS). Chromatin was reverse cross-linked overnight at 65°C in 0.2 M NaCl, followed by RNase A and proteinase K treatment, and the DNA purified using the ChIP DNA Clean & Concentrator kit (Zymo Research). ChIP DNA from two biological replicates were pooled and subjected to 2 x 150 bp paired-end read sequencing on the Illumina HiSeq-4000 platform (Genomics Resource Center, Institute for Genome Sciences, University of Maryland) to a sequencing depth of >14 million reads per sample. FANCD2 ChIP-seq datasets are accessible through GEO Series accession number GSE183048.

### ChIP-seq: Brd4 and H3K27ac

W12 chromatin samples were isolated as described above, and have been described previously ^25^. However, only the sequence data flanking the HPV16 integration sites was previously analyzed and published. Here, we analyze the same dataset but for the entire human genome. Antibodies used were Brd4 (Bethyl Laboratories A301-985A, 3 μg) or H3K27ac (Millipore 07–360, 3 μl). No antibody controls were included to monitor non-specific binding. Brd4 and H3K27ac ChIP-seq datasets are accessible through GEO Series accession number GSE183048.

### ChIP-seq processing and peak calling

Reads were trimmed with Cutadapt version 1.18 ^89^. All reads aligning to the ENCODE hg19 v1 blacklist regions ^90^ were identified by alignment with BWA version 0.7.17 ^91^ and removed with Picard SamToFastq, https://broadinstitute.github.io/picard/. Remaining reads were aligned to an hg19 reference genome using BWA. Reads with a mapQ score less than 6 were removed with SAMtools version 1.6 ^92^ and PCR duplicates were removed with Picard MarkDuplicates. Peaks were called by comparing each ChIP sample to its matching input sample. For FANCD2, the mean fragment size was estimated by Phantompeakqualtools version 2.0 ^93^. Peaks were called using SICER version 1.1 ^94^ with the following parameters: redundancy threshold of 100, effective genome fraction of 0.75, window size of 25,000 bp, and gap size of 50,000 bp. H3K27ac and Brd4 peaks were called using macsBroad (macs version 2.1.1 from 2016/03/09) ^95^ with the following parameters: --broad-cutoff 0.01 -f “BAMPE”. Data was converted into bigwigs for viewing and normalized by reads per genomic content (RPGC) using deepTools version 3.0.1^96^ using the following parameters: --binSize 25 --smoothLength 75 -- effectiveGenomeSize 2700000000 --centerReads --normalizeUsing RPGC. RPGC-normalized input values were subtracted from RPGC-normalized ChIP values of matching cell type genome-wide using Deeptools with --binSize 25.

### FANCD2-associated fragile site dataset

FANCD2 ChIP-seq peaks were filtered by a −log10 q-value of ten or above to remove low-confidence calls. Filtered C33-A ChIP-seq peaks were combined with previously mapped aphidicolin-induced FANCD2 peaks identified by ChIP-chip in these cells ^26^. C33-A (**Supplementary Table S8**) and HeLa FANCD2-enriched regions (**Supplementary Table S9**) were combined and overlapping peaks merged using bedtools MergeBED ^97^. Association of ChIP peaks between the three FANCD2 datasets were determined by permutation testing using regioneR ^98^. Combined FANCD2 peaks from C33-A and HeLa are listed in **Supplementary Table S10**. FANCD2-enriched regions were compared to aphidicolin-induced common fragile sites characterized in lymphoblast cells (FRA regions) and mitotic DNA synthesis regions characterized in HeLa cells ^41,42^ that are listed in **Supplementary Table S11** and **S12**, respectively, using regioneR ^98^. FRA-regions were downloaded from the HGNC (HUGO Gene Nomenclature Committee) database (https://www.genenames.org/download/custom/) on 08/27/2020 using advanced filtering: *gd_locus_type = ‘fragile site’*.

### Overlap analysis of FANCD2 enriched regions with long genes

The Gencode Release 19 human reference genome (GRCh37) was filtered for protein-coding genes greater than or equal to 0.3 Mb in length, including UTRs (**Supplementary Table S13**), and used to determine the overlap with FANCD2-enriched regions.

### Enhancer dataset

Consensus peak sets for H3K27ac were defined as overlapping regions found in at least four out of eight W12 samples using DiffBind ^99,100^. Enhancers were defined as genomic intervals that overlapped between H3K27ac peaks and Brd4 peaks and are listed in **Supplementary Table S14**. Proximal and distal enhancer-like cis-Regulatory Elements by ENCODE for NHEK and HeLa cells were downloaded from https://screen.encodeproject.org/<u>#</u> (accessed September 2020) ^51^. ENCODE GRCh38 enhancer files were converted to hg19 using the UCSC Genome Browser LiftOver tool, http://genome.ucsc.edu/cgi-bin/hgLiftOver. W12 enhancers were compared to ENCODE NHEK enhancers using regioneR ^98^.

### Overlap analysis of integration breakpoints with fragile sites and enhancers

The intersect between the genomic coordinates of HPV integration breakpoints (−/+ 50 Kb flank regions, ^7,9^) with enhancers (**Supplementary Table S14**) or FANCD2-enriched regions (**Supplementary Table S10**) was analyzed using the *Overlapping Pieces of Intervals* function in the Galaxy genomics platform (https://usegalaxy.org/). Samples with multiple reported breakpoints within the same chromosome were classified as a single integration locus if the 5’ and 3’ most breakpoints were within 3 Mb of each other. This 3 Mb cut-off was based on manual analysis of the distance between clustered breakpoints identified by WGS and/or hybrid-capture technologies for the CESC and HNSCC datasets. Each integration locus was assigned a unique integration ID so that the number of breakpoints per integration could be categorized as a cluster. Each integration breakpoint was analyzed independently for their association with the genomic feature of interest, as well as with adjacent HPV breakpoints, which were classified as belonging to the same integration locus/cluster. For significance testing, the data was permutated 10,000 times to create an expected distribution of the overlap between integration breakpoints and loci with FANCD2-enriched regions or enhancers using regioneR ^98^.

### Super-enhancer dataset

Super-enhancers were defined in the Brd4 and H3K27ac W12 ChIP-seq datasets using the Rank Ordering of Super-Enhancers (ROSE) tool, using default parameters ^53,54^. Enhancers defined in W12 cells (**Supplementary Table S14**) were used as the input list of enhancers. Super-enhancers were defined by Brd4 and/or H3K27ac consensus peaks that mapped in at least six out of twelve W12 samples for the 20831, 20862 and 20863 subclones and are listed in **Supplementary Table S16**. Super-enhancers mapped in the 20861 subclone were excluded as they were masked by the amplified viral-host derived super-enhancer-like element at the HPV integration site in these cells ^25^.

### Gene ontology analysis of W12 enhancers

W12 enhancers that overlapped with CESC integration breakpoints (−/+ 50 Kb flanks) were analyzed using the Genomic Regions Enrichment of Annotations Tool (GREAT) using default parameters, accessed July 2021, http://great.stanford.edu/public/html/. All enhancers profiled in W12 cells (**Supplementary Table S14**) were used as the input list of background regions.

## DATA AVAILABILITY

The data discussed in this publication have been deposited in NCBI’s Gene Expression Omnibus ^101^ and are accessible through GEO Series accession number GSE183048 (https://www.ncbi.nlm.nih.gov/geo/query/acc.cgi?acc= GSE183048).

## ACKNOWLEDGEMENTS

We thank Justin Lack (NIH/NCBR/NCI/NIAID/FNL) for advice on statistical analyses and Susan Huse (NIH/NCBR/NCI/NIAID/FNL) for preliminary analysis of enhancer-capture at integration sites. The results published here are in part based upon data generated by the TCGA Research Network, https://www.cancer.gov/tcga. This work was funded by the Intramural Research Program of NIAID, NIH grant number ZIA AI001223 LVD, and NIH grant number AI153156 (JMP). The funders had no role in study design, data collection and analysis, decision to publish, or preparation of the manuscript.

## AUTHOR CONTRIBUTIONS

AAM supervised the project. AAM and AW conceived and designed the study. AW compiled datasets and performed ChIP-seq experiments. TEM processed ChIP-seq data. TEM designed and advised on statistical analyses. AW and TEM performed statistical analysis. AAM, AW and TEM analyzed the data. JP and JK identified integration breakpoints from CESC and HNSCC TCGA datasets. AAM and AW wrote the manuscript. All authors discussed, critically revised, and approved the final version of the article for publication.

---

**Figure S1.**
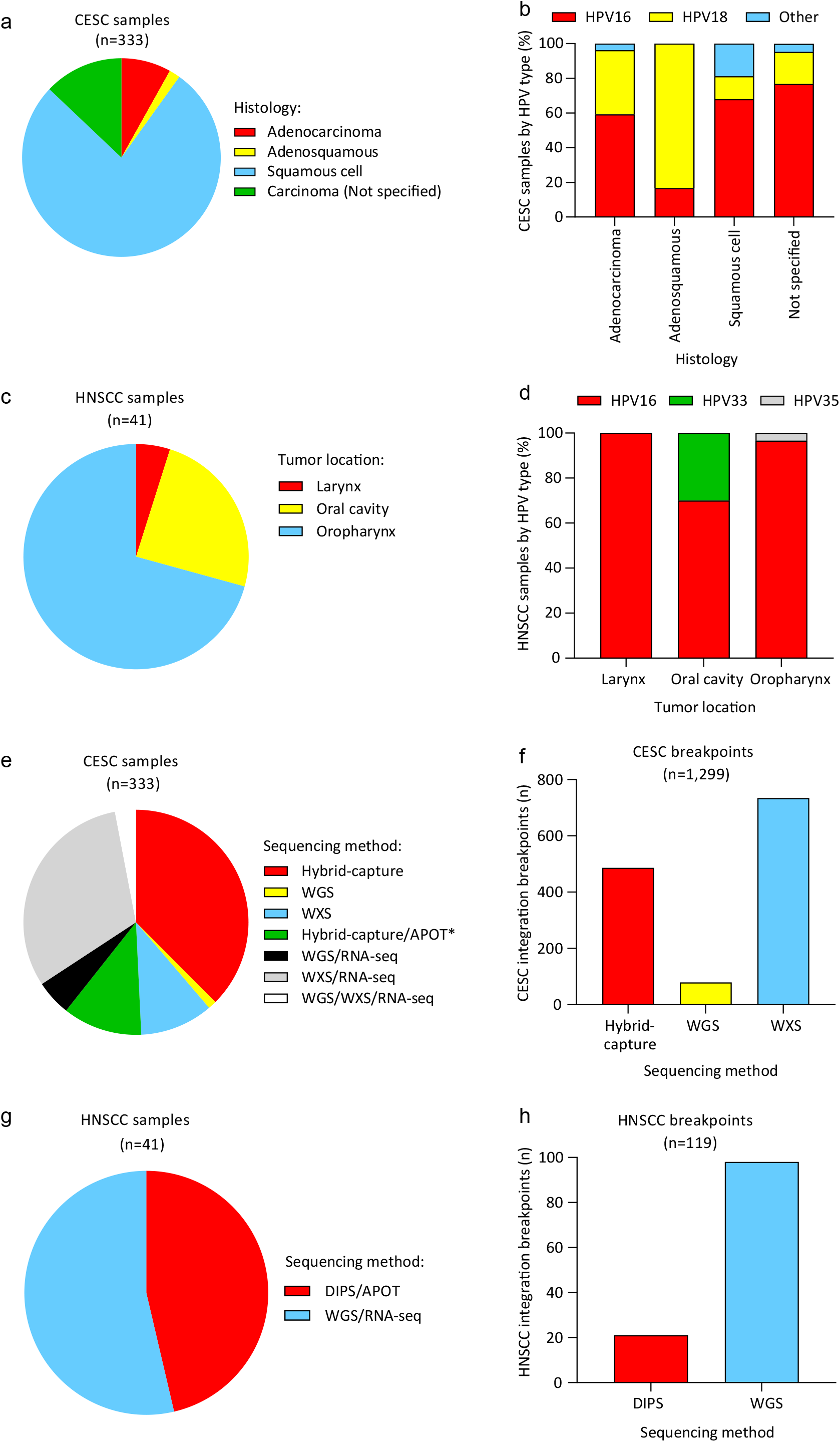
Characteristics of CESC and HNSCC HPV integration datasets. (**a**), The distribution of cervical carcinomas (CESC, n=333) based on tissue histology. Squamous cell carcinomas, adenocarcinomas and adenosquamous carcinomas accounted for 77.2%, 8.1% and 1.8% of cervical samples, respectively. CESC that were not specified by histology type accounted for 12.9% of the samples. (**b**), The frequency of HPV types across the different histological subtypes. One CESC sample (*TCGA-EA-A43B*) had two integration sites of different viral types and was therefore excluded from these counts. HPV16 (67.5%) and HPV18 (17.2%) were the most frequent HPV types detected in CESC. HPV16 was the predominant viral type in squamous cell carcinomas (HPV16, 68.0%; HPV18, 13.3%; other, 18.8%) and adenocarcinomas (HPV16, 59.3%; HPV18, 37.0%; other, 3.7%), whereas HPV18 was the predominant viral type in adenosquamous carcinomas (HPV18, 83.3%; HPV16, 16.7%; other, none). Head and neck squamous cell carcinomas (n=41) included in this study were predominantly male subjects (males, 82.9%; females, 17.1%), **Supplementary Table S2**. (**c**), Tumors of the oropharynx, including the base of tongue and tonsils, accounted for 70.7% of samples. The remainder of tumors were from the oral cavity (24.4%) and larynx (4.9%). (**d**), The percentage distribution of HPV type by HNSCC tumor location. HPV16 was the most predominant viral type in HNSCC (90.2%) and accounted for 96.6%, 70.0% and 100% tumors of the oropharynx, oral cavity and larynx, respectively. The remainder of HNSCC samples were positive for HPV33 (7.3%) and HPV35 (2.4%) that were isolated from the oral cavity and oropharynx, respectively. (**e-h**), The distribution of samples and integration breakpoints included in this study that were grouped by sequencing method. (**e-f**), For the CESC dataset, 50.8% samples had associated transcription-based data (**e**) and all integration breakpoints were detected through next-generation sequencing technologies (hybrid-capture/WGS/WXS with probes added to capture the integrated HPV DNA sequence) (**f**). (**g**), The HNSCC dataset was limited in size as >50% samples were identified from RNA sequencing alone and did not have matched DNA sequences. (**h**), Approximately 54% HNSCC samples were processed from WGS and 46% from DIPS, and all had matched RNA data (RNA-seq and APOT, respectively). *Two samples were also validated by RNA-seq analysis in addition to APOT. *APOT, Amplification of papillomavirus oncogene transcripts; DIPS, Detection of integrated papillomavirus sequences; WGS, Whole genome sequencing; WXS, Whole exome sequencing*.

**Figure S2.**
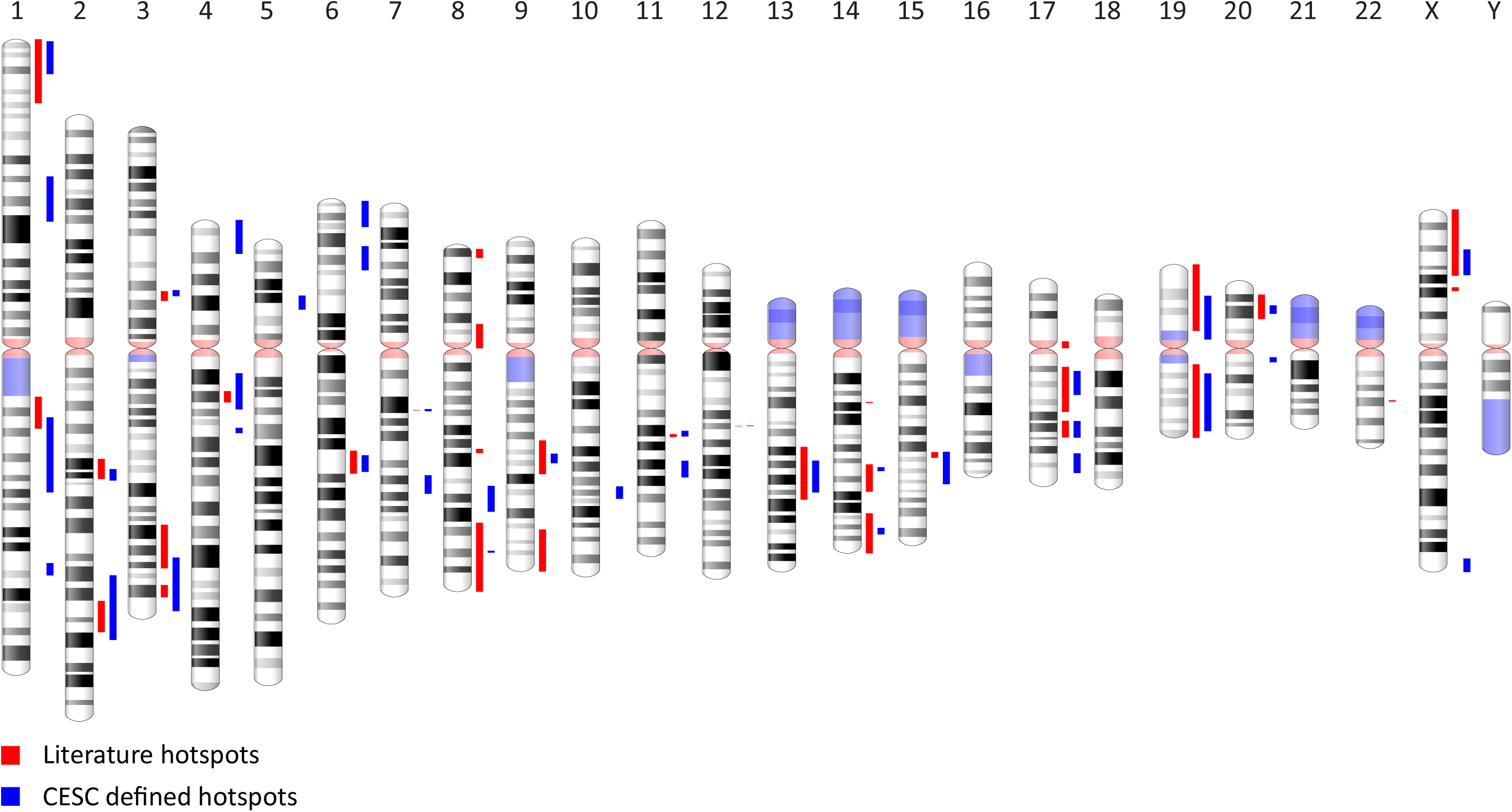
Integration hotspots defined in cervical tumors. Schematic representation of integration hotspots across the human genome defined previously in the literature (red bars) and in our cervical carcinoma, CESC, dataset (blue bars).

**Figure S3.**
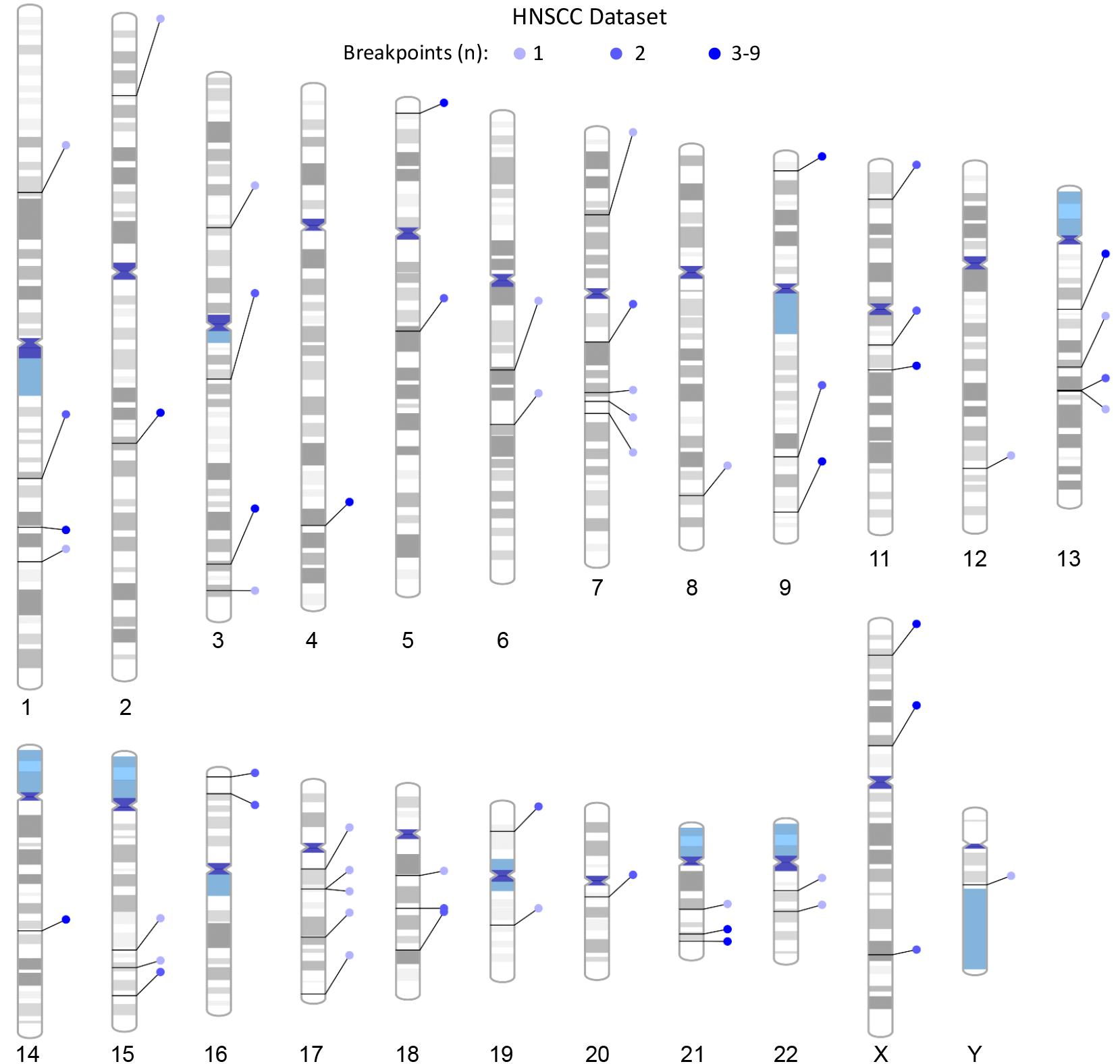
Distribution of clustered breakpoints at HNSCC integration loci. Schematic representation of clustered breakpoints at HNSCC integration loci across the human genome. Lines connecting to each chromosome represent different integration loci. Blue circles represent the indicated number of breakpoints per integration locus.

**Figure S4.**
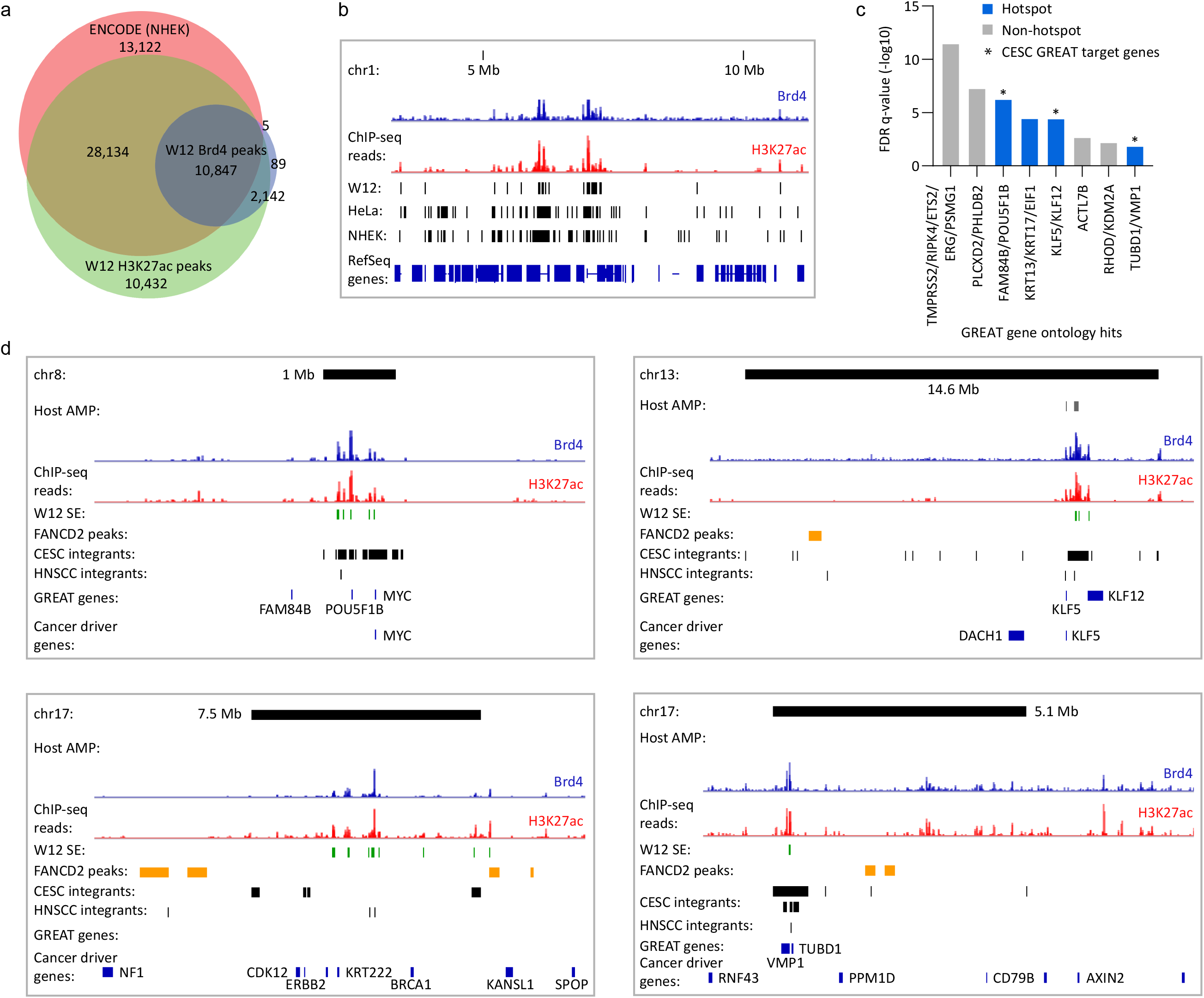
Keratinocyte-specific enhancers mapped by Brd4 and H3K27ac ChIP-seq. (**a**), Venn diagram showing the regions of overlap between Brd4 and H3K27ac ChIP-seq signals profiled in W12 cervical keratinocytes with enhancers defined in NHEK (Normal Human Epidermal Keratinocytes) cells from ENCODE ^51^. Permutation testing was used to determine the significance in overlap between W12 Brd4/H3K27ac consensus peaks and ENCODE NHEK enhancers (p<0.0001). (**b**), Alignment of Brd4 (blue) and H3K27ac (red) ChIP-seq signals mapped in W12 cervical keratinocytes with enhancers defined in the HeLa and NHEK ENCODE datasets. Black bars below the ChIP-seq signal tracks represent W12 enhancers that were defined by consensus peaks for Brd4 and H3K27ac enrichment. (**c**), GREAT (Genomic Regions Enrichment of Annotations Tool) gene ontology analysis was performed using W12 enhancers that overlapped HNSCC integration breakpoints (−/+ 50 Kb flanks) as input, and compared against all W12 enhancers, to identify putative target genes associated with these cis-regulatory regions. Bars represent putative target genes plotted against their FDR (false discovery rate) adjusted p-values (q-value). Blue and grey bars represent genes that overlap integration hotspots and sites of non-recurrent integration, respectively. Enriched target genes within the same genomic locus were grouped (e.g. KLF5 and KLF12) and plotted using the most significant q-value. (**d**), Alignment of Brd4 (blue) and H3K27ac (red) ChIP-seq signals mapped in W12 cervical keratinocytes at integration hotspots (black bars; size indicated in Mb) in cervical carcinomas. Grey bars represent amplified (AMP) host DNA in different HNSCC tumors from The Cancer Genome Atlas. Green, yellow, and black bars below the ChIP-seq signal tracks represent super-enhancers (SE) mapped in W12 subclones, FANCD2-associated fragile sites mapped in C33-A and HeLa cells and CESC/HNSCC integration loci, respectively. Genes identified from GREAT gene ontology analysis ^52^ and cancer driver genes ^55^ are indicated by blue bars.

**Figure S5.**
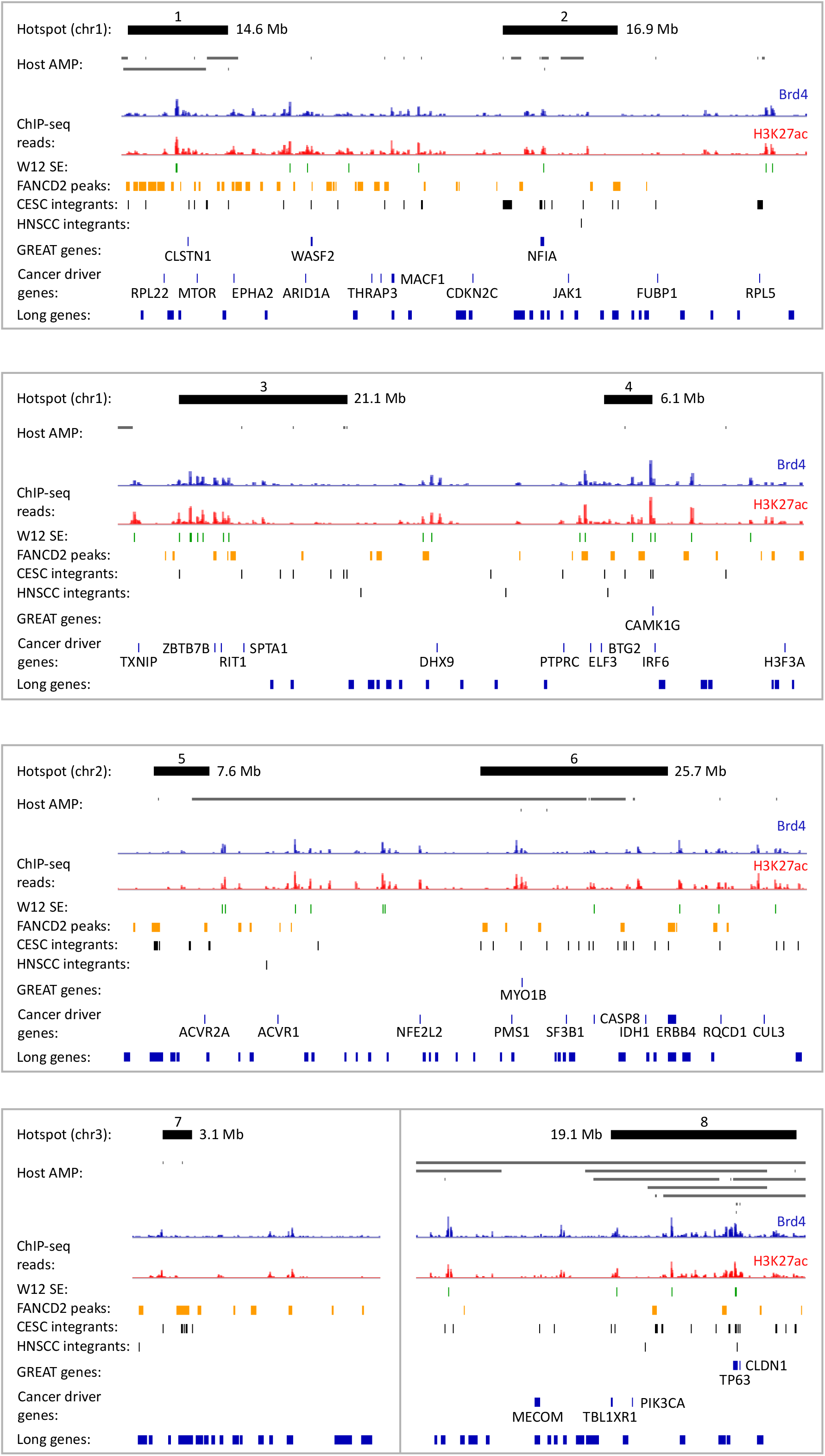
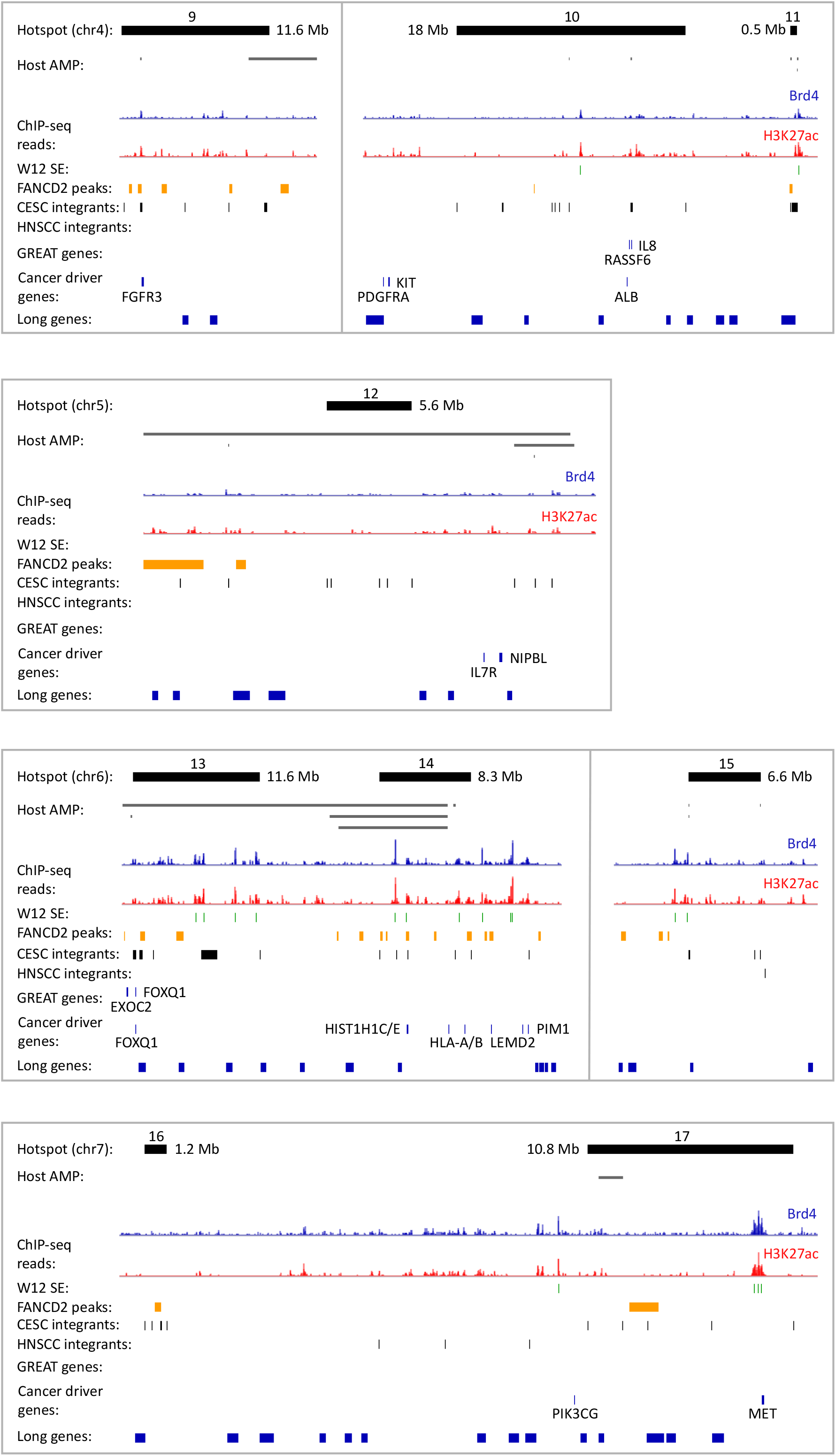
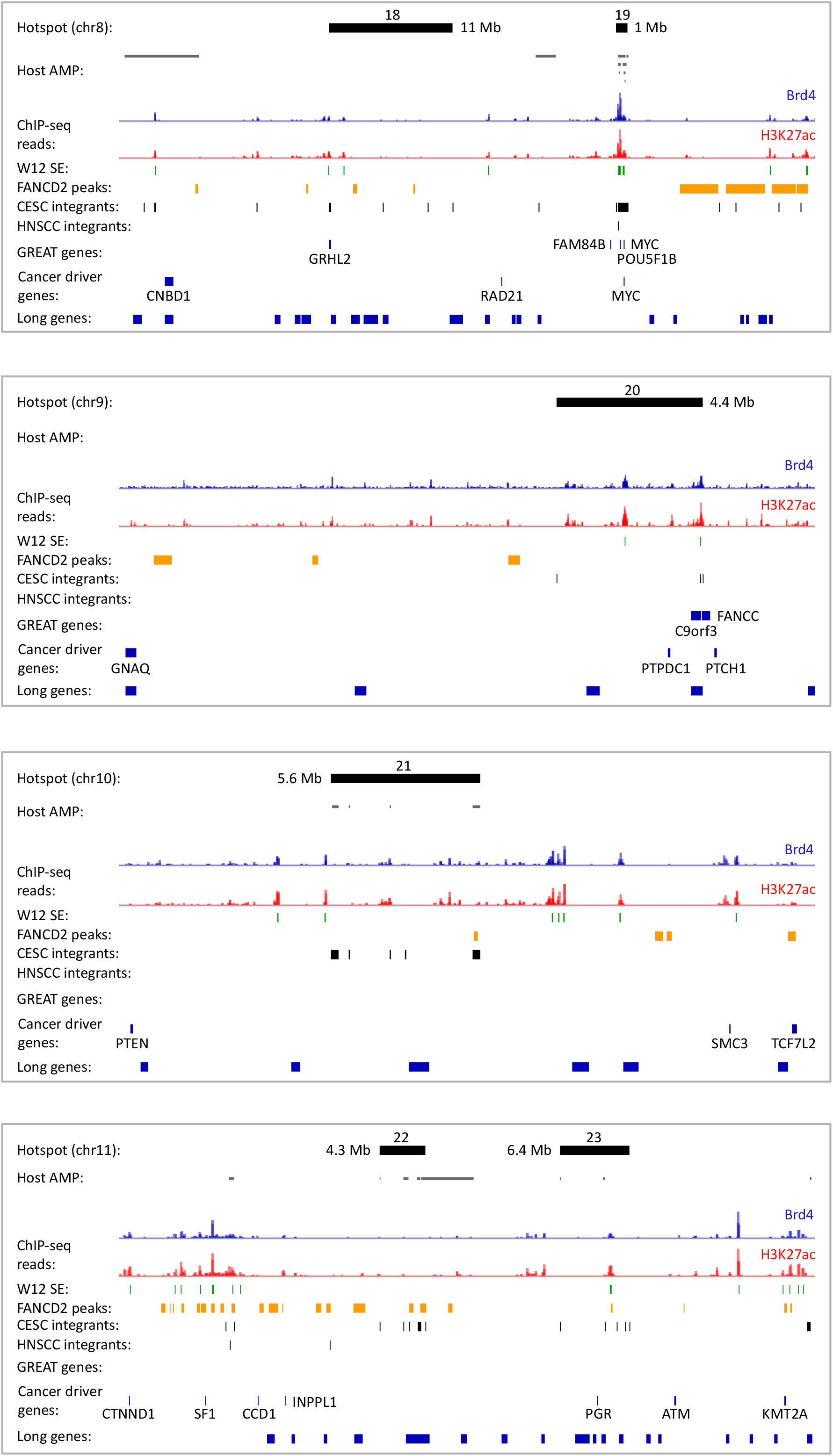
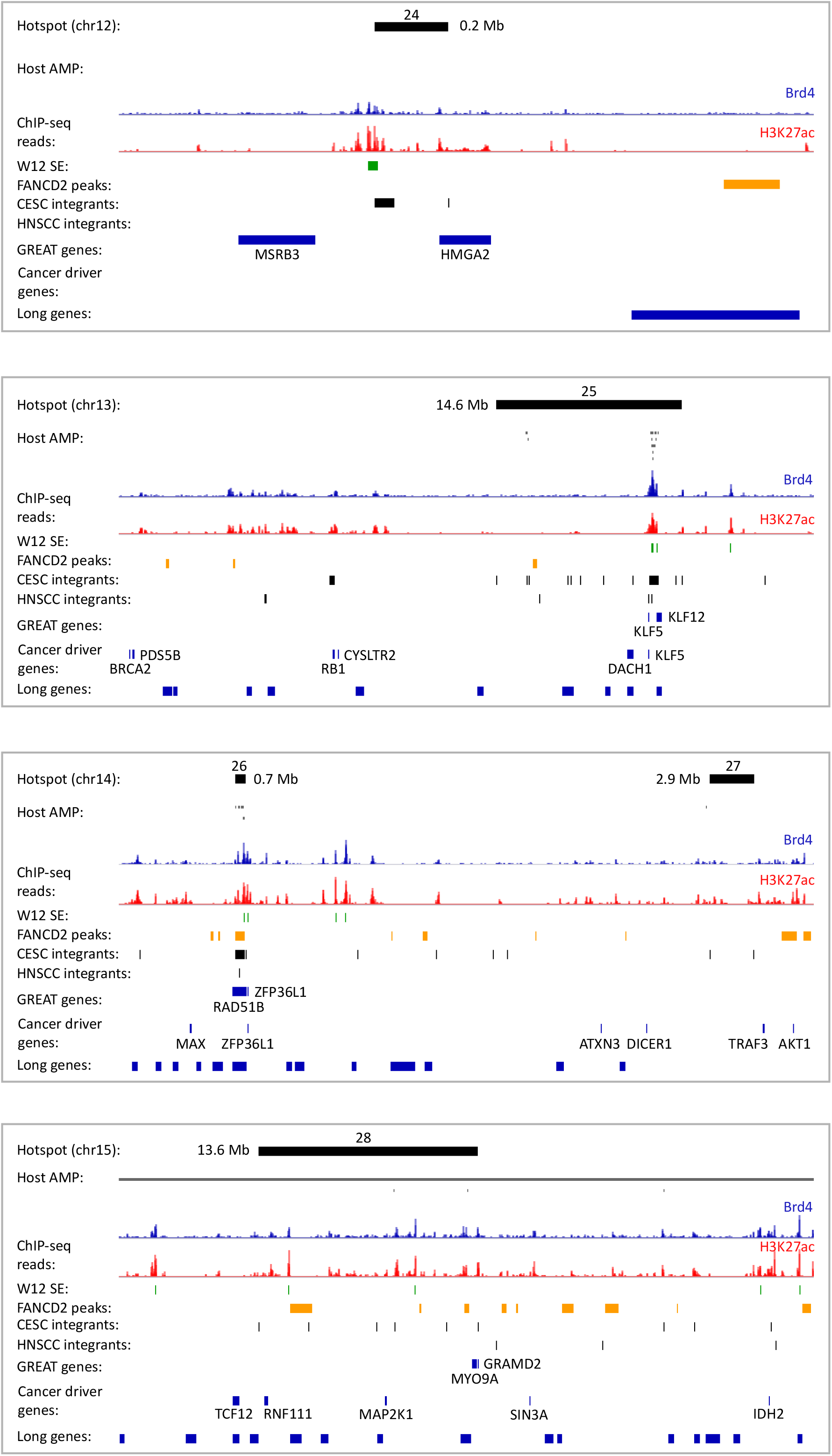
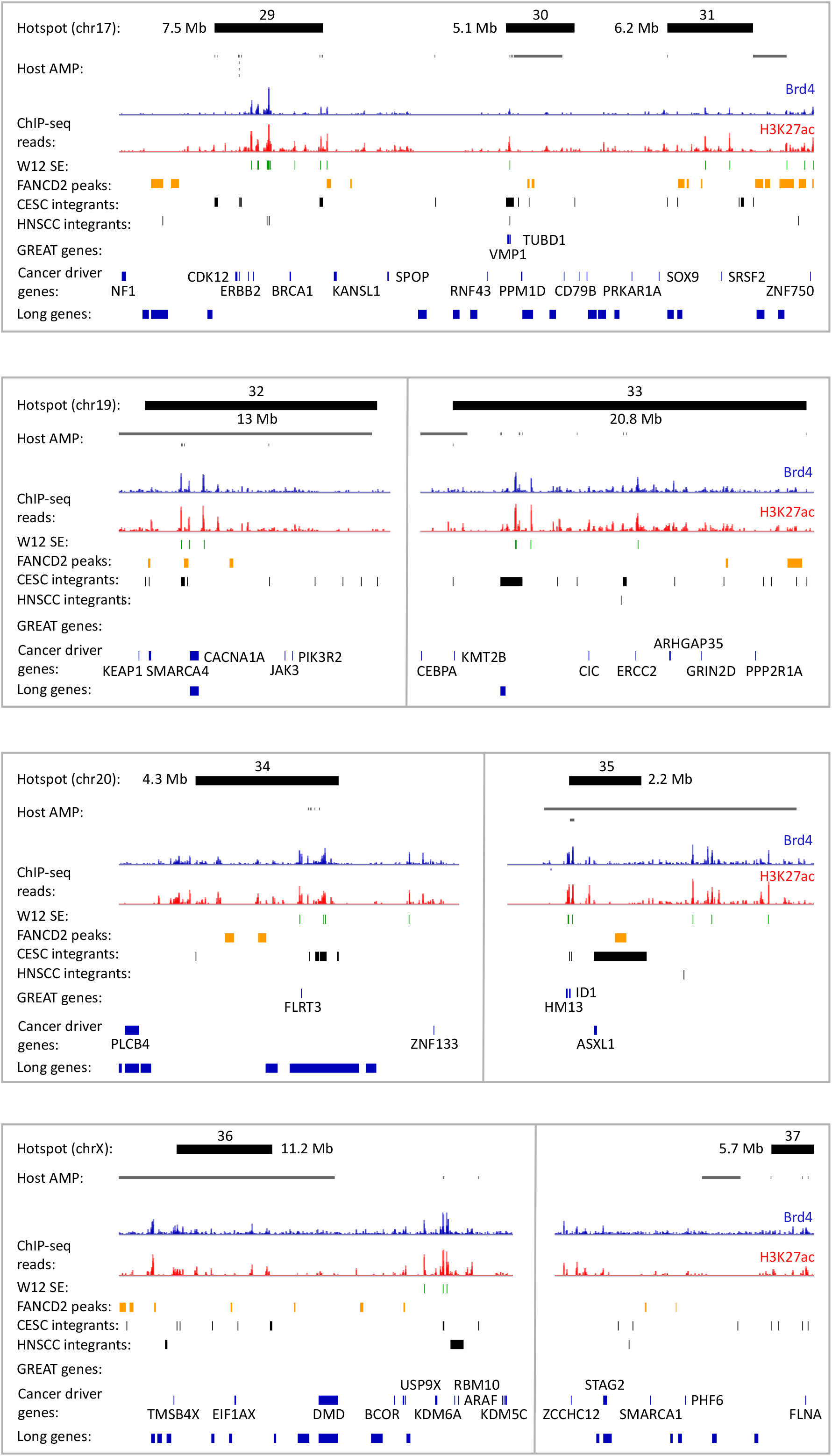
Genomic landscape of integration hotspots. Alignment of Brd4 (blue) and H3K27ac (red) ChIP-seq signals mapped in W12 cervical keratinocytes at integration hotspots (top black bars; size indicated in Mb) in cervical carcinomas. Numbers above each integration hotspot indicates the hotspot ID referenced in **Supplementary Table S4**. Grey bars represent amplified (AMP) host DNA in different CESC tumors from The Cancer Genome Atlas. Green, yellow, and black bars below the ChIP-seq signal tracks represent super-enhancers (SE) mapped in W12 subclones, FANCD2-associated fragile sites mapped in C33-A and HeLa cells and CESC/HNSCC integration loci, respectively. Long protein-coding genes (>0.3 Mb in length), genes identified from GREAT gene ontology analysis ^52^ and cancer driver genes ^55^ are indicated by blue bars. Each integration hotspot is further characterized in **Supplementary Table S4**.

## SUPPLEMENTARY TABLES

**Table S1. CESC and HNSCC datasets**

Cases represent the number (n) of patient samples with integrated HPV genomes detected by DNA and/or RNA sequencing methods. For hybrid-capture method, only integration breakpoints that were validated by Sanger sequencing were included in this study. Somatic copy number alteration (SCNA) datasets for CESC and HNSCC TCGA samples were downloaded from the Broad Institute ^86,87^.

**Table S2. CESC HPV integration breakpoints (n=1,299)**

CESC HPV integration sites (Patient ID, chromosome, integration breakpoint, HPV type, HPV breakpoints, tumour type, detection method used for calling integration breakpoints, target gene, (viral oncogene) transcription status, SCNA status, author of study, Pubmed ID). Only integration breakpoints detected through DNA sequencing were used for overlap analyses.

**Table S3. HNSCC HPV integration breakpoints (n=119) included in this study**

HNSCC HPV integration sites (Patient ID, chromosome, integration breakpoint, HPV type, HPV breakpoints, tumour location, detection method used for calling integration breakpoints, target gene, (viral oncogene) transcription status, SCNA status, gender, smoking status, author of study, Pubmed ID). Only integration breakpoints detected through DNA sequencing were used for overlap analyses.

**Table S4. Integration hotspots defined in CESC**

**Table S5. Integration hotspots defined in the literature**

Overlapping intervals for hotspots identified from the literature were merged into a single interval using the MergeBED tool (Merged hotspot position) for comparison to hotspots defined in our CESC dataset (**Supplementary Figure S2**).

**Table S6. CESC integrations with associated SCNA included in this study**

Samples that had associated host genome amplification data were classified based on the somatic copy number alteration (SCNA) status at the site of integration. Normal, normal genomic profile; AMP, amplification; DEL, deletion; DEL (excluded; overlapping deletion), deletions that overlap integration breakpoints were excluded as these likely reflect the alternative chromosome as sequencing of the chimeric viral-host junction was available for the associated HPV insertion sites

**Table S7. HNSCC integrations with associated SCNA included in this study**

Samples that had associated host genome amplification data were classified based on the somatic copy number alteration (SCNA) status at the site of integration. Normal, normal genomic profile; AMP, amplification; DEL, deletion; DEL (excluded; overlapping deletion), these likely reflect the alternative chromosome as sequencing of the chimeric viral-host junction was available for the associated HPV insertion sites

**Table S8. FANCD2-associated fragile sites mapped in C33-A cervical carcinoma cells**

C33-A FANCD2 peaks identified by ChIP-seq, filtered by −log10 q-value >10, combined with FANCD2 ChIP-chip peaks previously reported in C33-A cells ^26^.

**Table S9. FANCD2-associated fragile sites mapped in HeLa cervical carcinoma cells**

HeLa FANCD2 peaks identified by ChIP-seq, filtered by −log10 q-value >10.

**Table S10. FANCD2-associated fragile sites mapped in C33-A and HeLa cervical carcinoma cells**

C33-A FANCD2 peaks identified by ChIP-chip and ChIP-seq (**Supplementary Table S6**) were combined with HeLa ChIP-seq peaks (**Supplementary Table S7**). Overlapping FANCD2 peaks from the C33-A and HeLa datasets were merged (merging based on nearby peaks was set to 0 bp).

**Table S11. Aphidicolin-induced common fragile sites from the HGNC Database**

https://www.genenames.org/download/custom/. Bold font indicates FRA-regions that overlap with FANCD2-enriched regions listed in **Supplementary Table S8**. A total of 43/77 (55.8%) FRA-regions overlap with FANCD2-enriched regions profiled in C33-A and HeLa cells.

**Table S12. Mitotic DNA synthesis (MDS) regions that overlap with FANCD2 peaks**

MDS regions profiled in HeLa cells ^41,42^ were compiled and overlapping regions merged (merging based on nearby peaks was set to 0 bp). A total of 112/232 (48.3%) MDS regions overlapped with FANCD2-enriched regions profiled in C33-A and HeLa cells.

**Table S13. FANCD2-enriched regions that overlap long genes**

The Gencode Release 19 human reference genome (GRCh37) was filtered to identify protein-coding genes longer than or equal to 0.3 Mb, including UTRs. A total of 185/782 (23.7%) long genes overlapped with FANCD2-enriched regions profiled in C33-A and HeLa cells and are indicated in bold font. Genes that are transcriptionally active in C33-A and/or HeLa cells were determined from RNA-seq (^26^ and Expression Atlas, accessed September 2020, https://www.ebi.ac.uk/gxa/home) and are indicated in blue font. 121/185 (65.4%) long genes that overlapped with FANCD2-enriched regions are expressed in C33-A and/or HeLa cells.

**Table S14. Brd4/H3K27ac consensus enhancer peaks profiled in W12 subclones**

Brd4 and H3K27ac peaks were identified through ChIP-seq in HPV16-positive W12 cervical keratinocyte subclones (20831, 20861, 20862 and 20863). Consensus peaks for H3K27ac were identified across the four W12 subclones. Enhancers were defined as the overlapping genomic intervals between H3K27ac consensus peaks and Brd4 peaks.

**Table S15. Top 20 biological processes associated with enhancers that overlap integration breakpoints in CESC and HNSCC**

**Tables S16. Super-enhancers defined in W12 cervical keratinocytes**

**Table S17. Cancer driver genes within 1 Mb of super-enhancers that overlap integration loci**

